# Dynamic control of visually-guided locomotion through cortico-subthalamic projections

**DOI:** 10.1101/2020.02.05.936443

**Authors:** Elie M. Adam, Taylor Johns, Mriganka Sur

## Abstract

Goal-directed locomotion requires control signals that propagate from higher-order areas to regulate spinal mechanisms. The cortico-subthalamic hyperdirect pathway offers a short route for cortical information to reach locomotor centers in the brainstem. We developed a task where head-fixed mice run to a visual landmark then stop and wait to collect reward, and examined the role of secondary motor cortex (M2) projections to the subthalamic nucleus (STN) in controlling locomotion. Our modeled behavioral strategy indicates a switching point in behavior or a sudden change from running to stopping, suggesting a critical neuronal control signal at stop locations. Optogenetic activation of M2 axons in STN leads the animal to stop prematurely. By imaging M2 neurons projecting to STN, we find neurons that are active at the onset of stops when executed at the landmark but not when executed spontaneously elsewhere. Our results suggest that the M2-STN pathway can be recruited during visually-guided locomotion to rapidly and precisely control the mesencephalic locomotor region (MLR) through the basal ganglia. By capturing the physiological dynamics through a feedback control model and analyzing neuronal signals in M2, MLR and STN, we find that the cortico-subthalamic projections potentially control MLR activity by differentiating an M2 error signal to ensure fast input-output dynamics.

## Introduction

Coordinated movement, and in particular locomotion, is enabled by distributed spinal and brain circuits. Although the executive mechanisms for locomotion are implemented in the spinal cord (Goulding, 2009; Grillner, 2003; Kiehn, 2016, 2006), locomotion is importantly regulated by supraspinal circuitry (Ferreira-Pinto et al., 2018; Kim et al., 2017; Ryczko and Dubuc, 2013). Brainstem circuits can induce locomotion upon targeted stimulation (Caggiano et al., 2018; Capelli et al., 2017; Josset et al., 2018; Roseberry et al., 2016); however, goal-directed locomotion, especially when guided by sensory information, requires control signals from higher-order areas directed towards the spinal cord (Arber and Costa, 2018; Drew et al., 2004; Grillner et al., 2008). The repertoire of higher-order signals required to regulate locomotion, and the neural circuitry that enables such signaling, are particularly unclear in behavioral settings (Arber and Costa, 2018). The control principles and substrates governing action can further suggest principles for cognitive control, particularly if the same neural substrate supports multiple functions.

While the regulation of locomotion is mainly implemented in the brainstem, movement planning is considered to arise at the level of the cortex (Churchland et al., 2010; Economo et al., 2018; Shenoy et al., 2013; Svoboda and Li, 2018; Wong et al., 2015). The basal ganglia are situated between these two centers and importantly regulate voluntary movement (Bolam et al., 2000; Graybiel, 2000). The direct and indirect pathway, initiated from the striatum, bidirectionally control locomotion (Roseberry et al., 2016) through the substantia nigra pars reticulata (SNr) and its control over the mesencephalic locomotor region (MLR) (Freeze et al., 2013; Liu et al., 2018). The basal ganglia additionally admit a cortical input straight to the subthalamic nucleus (STN) (Nambu et al., 2002, 2000), which projects to SNr (Hamani et al., 2004). This pathway has been termed the hyperdirect pathway, and evidence across species, notably humans, has shown stop activity at the source of this pathway in reactive stop-signal or go/no-go tasks (Aron et al., 2016; Aron and Poldrack, 2006; Eagle et al., 2008; Wessel and Aron, 2017; Chen et al., 2020). The STN itself is considered pivotal in stopping movement (Hamani et al., 2004; Schmidt et al., 2013). In rodents, reducing STN’s excitatory output induces hyperlocomotion (Schweizer et al., 2014), and lesions of STN induce impulsive responding (Baunez and Robbins, 1997; Eagle et al., 2008; Uslaner and Robinson, 2006). More recently, optogenetic studies in mice show that areal activation of STN excitatory cells disrupts self-initiated bouts of licking (Fife et al., 2017) and that activation and inactivation of STN-projecting prefrontal cortex neurons reduced and increased inappropriate licking (Li et al., 2020). Thus, the hyperdirect pathway stands as an important short-latency cortico-brainstem route for fast control of locomotion.

Goal-directed locomotion implies a proactive locomotor plan that is implemented to achieve a needed goal. From an engineering standpoint, we can enforce a desired goal trajectory in a system through feedback control (Aström and Murray, 2010). Feedback control is based on an error signal: a measured discrepancy between what we would like the system to do (reference) and what it is actually doing (output). By processing such a signal through a controller and feeding it to the system (plant) we intend to control, we ensure adequate performance (Dahleh et al., 2004; Oppenheim et al., 1996). If such a principle is implemented in neural circuits, then we would expect surges in neural signals upon sudden changes in planned locomotion trajectories, and that such signals would drive movement corrections to ensure fast control.

We thus developed a task where head-fixed mice run to a visual landmark, then stop and wait to collect reward, and examined the projections from secondary motor cortex (M2) to STN. We hypothesized that these projections send rapid signals that halt locomotion. Here, we report the existence of such signals sent from M2 to STN that halt visually-guided locomotion. This positions the hyperdirect pathway as a controller onto the mesencephalic locomotor region (MLR) in the midbrain. Furthermore, using dynamical systems and control engineering methods, we find that the hyperdirect pathway potentially controls the MLR by differentiating an M2 error signal to ensure fast input-output dynamics.

## Results

### Mice were trained to run, stop and wait at visual landmarks to collect reward

We developed a task that allowed us to examine proactive visually-guided locomotion stops (**Figure 1A**). A head-fixed mouse was positioned on a self-propelled treadmill in a virtual runway flanked on both sides by a continuous streak of LEDs. At the start of a trial, the animal was presented with a visual landmark consisting of a lit contiguous subset of LEDs, at a variable position from the animal. The movement of the treadmill was coupled to the movement of the landmark; as the mouse rotated the treadmill to move forward, the landmark approached the mouse. The mouse was then required to run and stop at the landmark, holding its position for 1.5s to collect reward. If the mouse waited at the landmark for the required time, it received a reward tone and a water reward simultaneously. If the mouse either ran to the end of the runway, bypassing the landmark, or failed to stop at the landmark within 30s, it received a miss tone. After the reward or miss tone, all the LEDs were turned off and a new trial started after 1s, with a landmark reappearing (**Video S1**).

**Figure 1:**
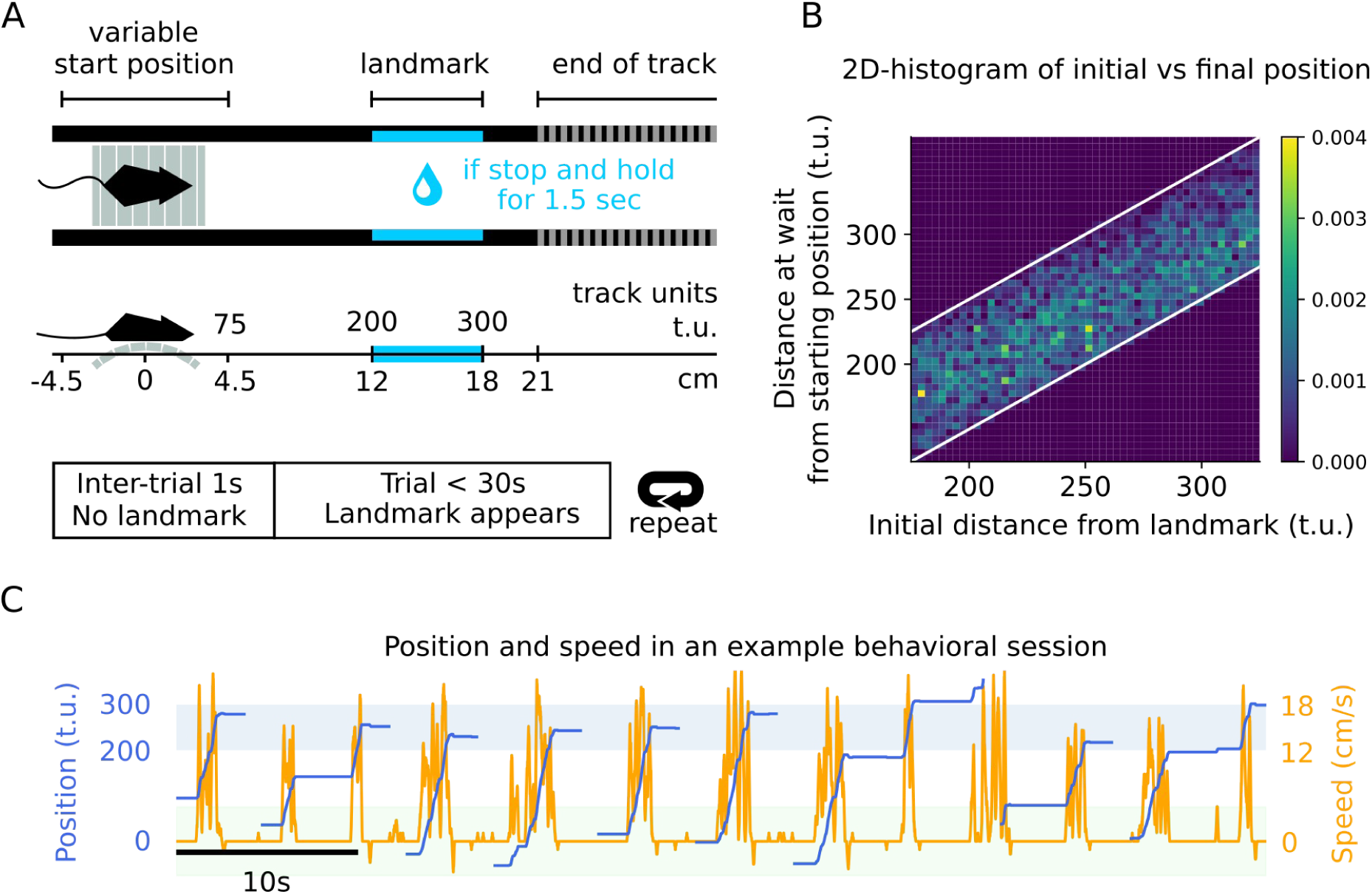
Mice were trained to run, stop and wait at visual landmarks to collect reward. **(A)** Schematic showing the task design. Position is defined in terms of track units (t.u.) with 200t.u. corresponding to 12cm. **(B)** Graph showing the distance at wait from starting position versus the distance from the landmark at the beginning of each trial. The color gradient indicates the frequency of stopping at the corresponding distance. The white lines indicate the beginning and end of the landmark. The distance of the landmark from the initial position does not affect the animal’s final stop position, indicating that the animals are using the visual cues, instead of relying on other mechanisms. N=10 mice, 3 sessions each, between 100-250 hit trials each. **(C)** Example data showing a mouse’s position on the track and its underlying speed for 9 trials. The blue and green shaded area indicates landmark position and potential starting position, respectively.

We ensured that the distance over which the landmark position randomly varied was greater than the width of the landmark. This prevented the animal from relying on tracking the distance to the landmark internally, ensuring that the task was, indeed, visually-guided (**Figures 1B and S1A,B**). The task elicited an ON-OFF locomotion pattern (**Figures 1C and S1A**), of which we carefully examined the stops. Furthermore, this pattern was a learned behavior (Figure S1F-I) as evidenced by the increase in stop-wait time as training sessions progress (Figure S1F) and an increase in hit-rates in a simpler version of the task during the first stages of training where landmarks are not present (Figure S1G).

### The behavior suggests a sudden switch in locomotion state

We modeled the behavior of the animal in a single trial as an optimal-control problem **(Figure 2A, Text S1)**. Starting from an initial position away from the landmark, the mouse is tasked to pick a locomotor plan (control policy) that dictates its locomotion pattern so as to minimize time to collect reward, thereby maximizing reward in a session. The notion of a trajectory guiding movement appears in various forms in the literature on the control of movement (McNamee and Wolpert, 2019; Shenoy et al., 2013; Wolpert and Ghahramani, 2000), and sets a basis for optimal feedback control in motor coordination (Todorov, 2004; Todorov and Jordan, 2002). We considered a simple setting, and modeled the dynamics relating the locomotion plan u_t_ (in track units/s) to the speed of the animal v_t_ by a first-order ordinary differential equation parametrized by a time-constant τ:

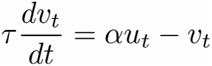

**Figure 2:**
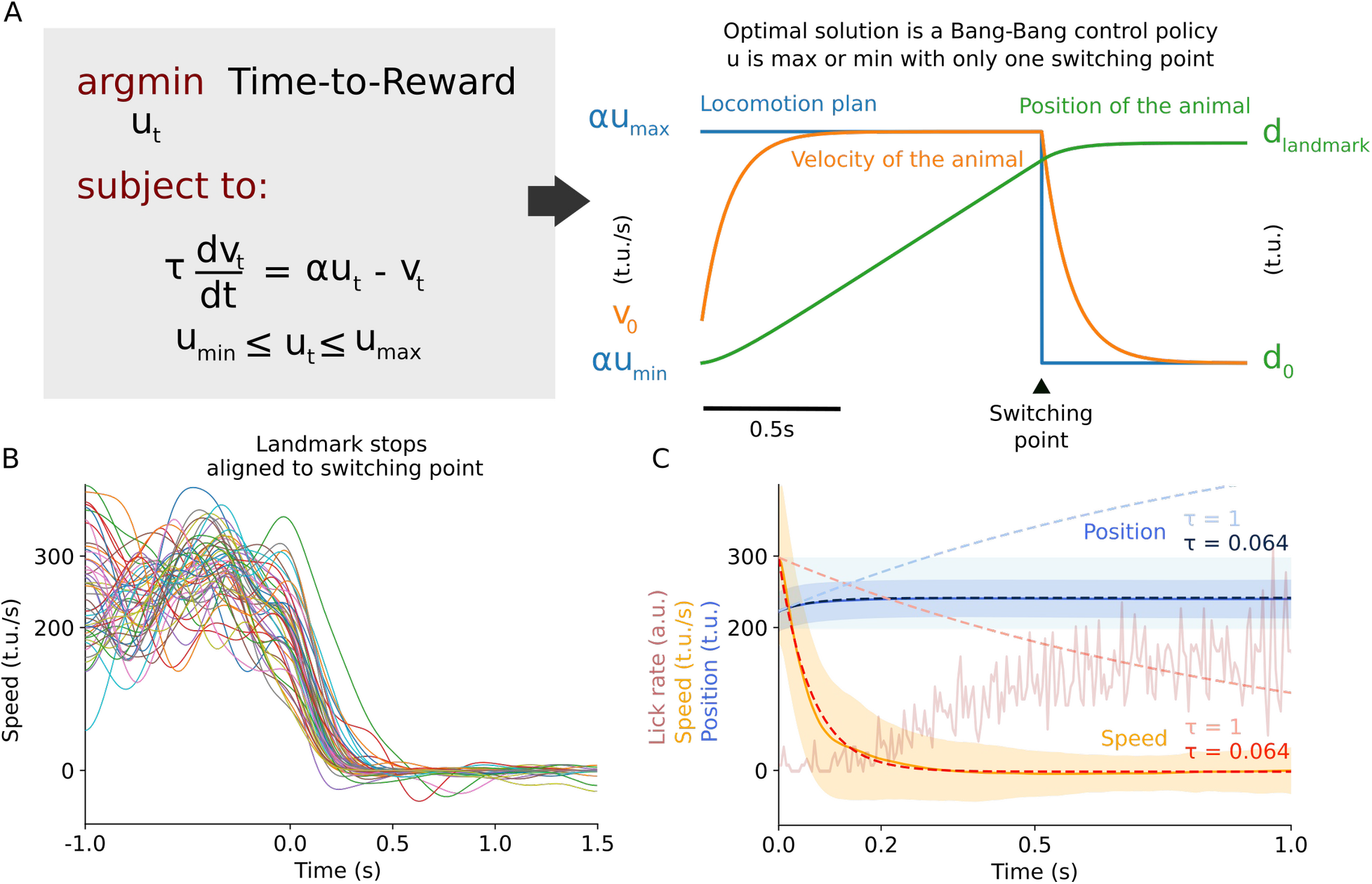
The behavior suggests a sudden switch in locomotion state. **(A) Left.** Formulation of the behavior as a minimum-time optimal-control problem. The mouse is tasked to pick a locomotor plan (u_t_) that minimizes the time required to collect reward. The locomotor plan dictates the speed of the animal, and the relation is governed by a first-order ordinary differential equation, parametrized by a time-constant τ. Furthermore, the locomotor plan, and therefore the speed, are bounded and cannot be infinite. **Right.** The optimal solution is a bang-bang control policy, where u_t_ starts at its maximum and suddenly switches to its minimum. **(B)** Examples of speed traces in landmark-stop windows aligned to the last peak in velocity before stopping indicating the ‘switching point’ (30 trials drawn from all the sessions across animals. N=10 mice, 3 sessions each). **(C)** Graph of the average speed, position, lick rate and model fits (dashed line) to actual data (continuous line) for speed and position in the period immediately following the switching point. Modeled position and speed using the equation in (A) identified a time constant τ = 63.75 ms are depicted with dark blue and red dashed lines. Predicted position and speed with τ = 1 s are depicted with light blue and light red dashed lines. The shaded regions correspond to the standard deviation of the sampled distribution. N=10 mice, 3 sessions each.

The parameter τ is fixed and dictates how quickly the speed follows the locomotion plan. The larger the value of τ, the less sensitive the change in velocity is to the discrepancy between the velocity and the locomotion plan. The scalar α makes our modeling more general (and has a default value of 1). We further bounded u_t_ by forcing it to be between two values u_min_ and u_max,_ thereby ensuring that the animal cannot run infinitely fast **(Figure 2A, Text S1)**. We then solved this control problem for the optimal solution via Lagrangian **(methods, Text S1)** and found it to be a bang-bang control policy: the optimal solution consists of having the locomotor plan be at u_max_ from the start of the trial up to a switching time, where it abruptly changes to u_min_. Many trajectories satisfy the constraints of the problem and are solutions, particularly ones where the animal slowly ramps down its speed as it approaches the landmark to stop. However, these strategies yield a longer time to reward, and the best (optimal) one, adopted by the animal if it aims to minimize time to reward, is one where the change is abrupt.

Therefore, to collect reward as soon as possible, the animal should accelerate as much as possible up to a switching point, which likely occurs before the animal arrives at the landmark, then suddenly brake as much as possible to arrive at a full halt at the landmark. This model yields two features. First, the optimal solution suggests that there is an essential switching point in behavior: if the brain generates a signal to stop, it should occur around this switching point. Second, by changing the value of τ, we can modulate how quickly the animal stops. Signatures of these strategies were later analyzed in the neurophysiological recordings.

From each session, we recovered time windows around all the stops at the landmark, which we termed landmark-stop windows, and we let the switching point in each correspond to the time point where the speed of the animal last peaked before stopping. We then aligned all the speed trajectories in the time windows along their switching point (**Figure 2B**). The model fitted well to the average trajectory with an average time constant of 63.75ms (**Figures 2C and S1C-E**) (**Methods**). Importantly, having a small enough time constant was essential for the animal to stop in a timely manner at the landmark. For instance, if τ were equal to 1s, the animal would not stop in time and would miss the landmark (**Figure 2C**). [The velocity fluctuated while the animals were running, leading to multiple peaks that were not followed by full stops. The model is not designed to capture these fluctuations, but to capture the decay speed during landmark stops which is independent of the speed profile before stopping. Indeed, the physical velocity engages many downstream brainstem and spinal circuits and is subject to locomotor contraints that forces it to be noisy and fluctuating around the theoretical speed. Of course, such fluctuations are not observed when the animal is not running, and the rate of decay does not depend on the speed amplitude, reflecting the transition from the ON state to the OFF state independently of the fluctuations.]

Our task was designed to elicit short bouts of runs and stops, instead of lengthy periods of locomotion (Methods). Our task design was intended to elicit locomotion in the ‘moderate velocity’ regime (as opposed to high velocity behaviors such as escape behavior) with the goal of engaging the pedunculopontine nucleus (PPN) in the MLR, and more likely basal ganglia circuitry, instead of the cuneiform nucleus (Caggiano et al., 2018; Josset et al., 2018). Importantly, the trajectory of the speed signal was not affected by the distance of the landmark **(Figure S1J)** if kept in close rang**e.** In all subsequent experiments, the trajectory of the stop was preserved throughout **(Figure S1K)**.

### Activating M2 axons in STN leads to stopping

Our behavioral model indicates a switching point in behavior, and suggests that the brain generates a signal at that time. As we expect the signaling to be the result of an instantiated locomotor plan, frontal associative regions are good candidates to contain such plans. In fact, activity emanating from the right inferior frontal cortex and the pre-supplementary motor area in humans has generally preceded stops in stop-signal reaction tasks and go/no-go tasks (Aron and Poldrack, 2006; Eagle et al., 2008; Nachev et al., 2008; Swann et al., 2012). The pre-supplementary motor area in humans is of particular interest. Its homologous structure in mice is unclear; however, some of its properties have been often considered to coincide with the medial part of M2 (Barthas and Kwan, 2017). Additionally, the task is visually guided, and medial M2 has been considered to be part of the visuomotor subnetwork in the mouse brain (Barthas and Kwan, 2017; Zhang et al., 2016, 2014; Yamawaki et al., 2016). STN receives projections from most of the frontal cortex, and medial M2 is linked with the associative subdivision of the BG and STN, which may have a role in visuomotor transformation and integration (Alexander et al., 1986; Bolam et al., 2000; Hamani et al., 2004; Hintiryan et al., 2016; Hooks et al., 2018; Mandelbaum et al., 2019). We hypothesized that the M2-STN pathway may offer a quick route for information to achieve rapid stops. Thus, we asked: if activity is sent along the M2-STN pathway, does it lead the animal to stop locomoting?

To address this question, we injected an adeno-associated virus (AAV) expressing ChR2 under the CaMKII promoter in M2, and implanted an optic-fiber above ipsilateral STN to target the M2 axons there (**Figure 3A, B**). On a random subset of trials, we delivered a brief burst of blue (473nm) light (at 20Hz for 500ms) into the optic-fiber to activate the axons once the animal crossed the middle of the runway, while running towards the landmark (**Figure 3A**). We found that activating the axons led the animal to stop prematurely (**Figure 3C-F**). This suggests that if the brain transmits a signal along the hyperdirect pathway right before stopping, it will causally trigger the animal to stop. However, the optogenetic activation was not effective at inducing premature stopping in all trials (**Figure 3F**), likely due to it being unilateral and to the presence of redundant circuits whose activity alteration might be necessary to initiate stopping. To verify that this effect was not due to inadvertedly activating fibers beneath STN in the cerebral peduncle (and not only M2 boutons in STN) because of spurious laser effects reaching that area, we instead targeted the M2 efferents anterior to the STN by placing the optic fiber above the cerebral peduncle, we found that stimulation did not affect locomotion (**Figure S2A-C**). Furthermore, when injected an AAV expressing only GFP (**Figure S2D,E**) and placing an optic fiber above STN, we found no premature stopping during the optogenetics control experiment upon laser delivery (**Figure S2F)**.

**Figure 3:**
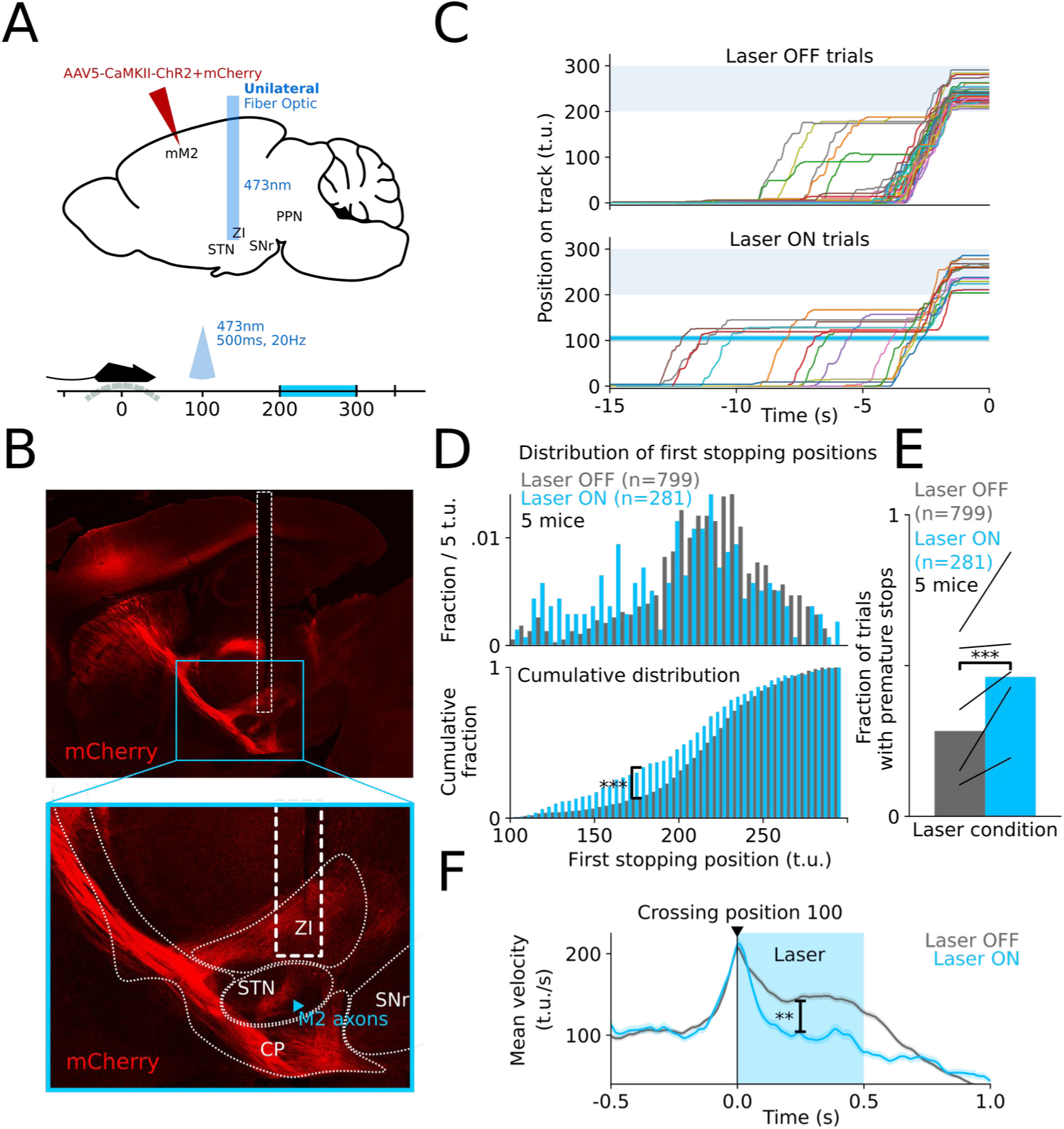
Activating M2 axons in STN leads to stopping. **(A)** An AAV virus expressing ChR2 was unilaterally injected in M2 of wild-type mice (N=5) and an optic fiber implanted over STN (ipsilateral to the injection site) to optogenetically target M2 axons in STN. On pseudorandom trials, blue light (473nm) was presented for 500ms at 20Hz as soon as the animal crossed track position 100. **(B)** M2 projects directly to STN via the hyperdirect pathway. The image shows the projections (mCherry) and a fiber-optic placement. **(C)** Position traces showing laser ON and OFF hit trials aligned to reward time for 1 session in 1 animal. The blue line indicates the position at which the laser is turned ON. To ensure enough running distance to position 100 and have it be a mid-point, the optogenetics sessions were performed at a fixed starting position of 0, although mice were trained on variable landmark distance. **(D)** Plots showing the distribution of the first position the animal stops at after position 100. We observe a shift in the distribution, towards position 100, indicating that during laser ON trials the animal stopped prematurely (Kolmogorov–Smirnov test, ***:p=4.76e-8<0.005). **(E)** Plot showing the fraction of hits trials with premature stops (stopping first before position 200) during laser ON and OFF trials (Mann-Whitney U test, ***:p=0.0<0.005). **(F)** Plot showing the average speed of the animal in Laser ON and OFF trials aligned to the time of crossing position 100, with a significant difference between Laser ON and OFF trials after crossing position 100 (Mann–Whitney U test, **:p=0.0034). The blue shaded area shows the period in which the laser is delivered. The peak of velocity at position 100 is an effect from averaging: while speed fluctuates, it is high while the animal is crossing position 100. Averaging will then yield a high average speed at position 100, and lower average elsewhere where peaks and troughs in speed are averaged.

### Stop activity is seen in M2-STN neurons on landmark stops but not mid-stops

We next asked: is there activity at the onset of stopping in the M2-STN pathway? To answer this, we imaged the calcium activity (GCaMP6f) of M2 neurons projecting to STN using two-photon microscopy. For each imaged neuron, we retrieved the activity during the landmark-stop windows (1s before the switching point and 1.5s after), averaged it across these windows and found a range of responses in different epochs (**Figure 4A**).

**Figure 4:**
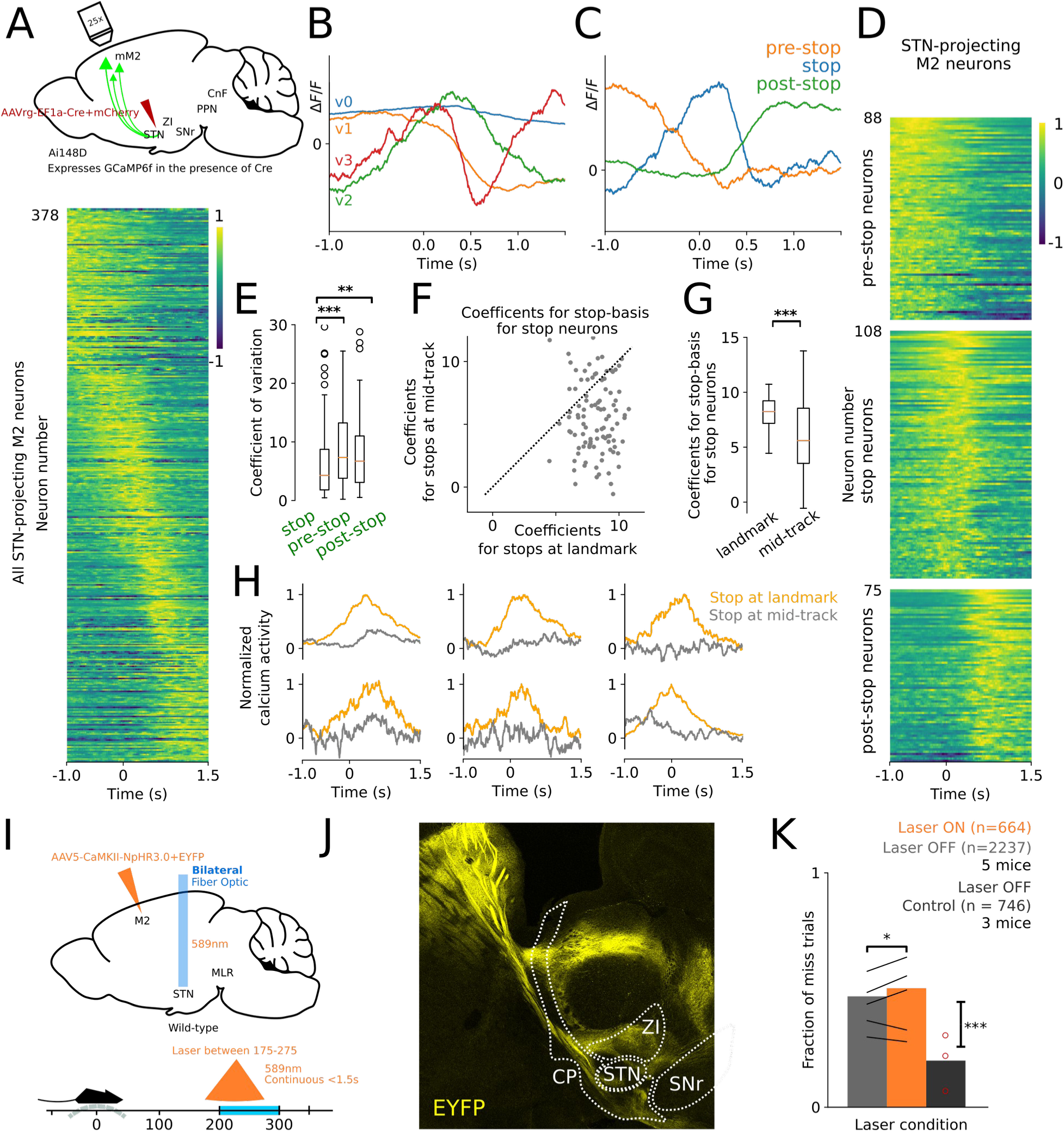
Stop activity is seen in STN-projecting M2 neurons at landmark stops but not at mid-stops. **(A) Top.** An AAV virus was injected in STN of Ai148D mice (N=4) to retrogradely express Cre, and thereby GCaMP6f in M2 neurons projecting there. **Bottom.** Plot showing the normalized average calcium activity of labeled M2 neurons within landmark-stop windows aligned to the switching point. **(B)** Graph showing a basis for a four-dimensional subspace that captures more than 70% of the energy in the responses, all pooled together. **(C)** Graph showing 3 templates of ideal neuronal responses, derived through a change of basis from the templates in (B), that are selectively active in different epochs in the task: before stops (pre-stop), during stops (stop) and after stops (post-stop). Each neuronal response can then be expressed as a weighted combination of these 3 templates. **(D)** Plot showing neurons (n=271), whose neural response energy (area under the squared signal) was more than 80% explained by the subspace, clustered into three groups (pre-stop, stop and post-stop) using the templates in (B). **(E)** Boxplots showing the reliability of neuronal responses during the stop period for each of the three clustered population in (D). High reliability is measured by a low coefficient of variation. Stop neurons show a significantly higher reliability compared to pre-stop (Mann-Whitney U test, ***:p=6.3e-4<0.005) and post-stop neurons (Mann-Whitney U test, *:p=0.011<0.05). **(F)** Scatterplot of the coefficient of contribution of the stop template in (C) in spontaneous-stops versus landmark-stops. Each data point represents a stop neuron (N = 108) taken from (D), and the coefficient represents the energy of the response along the dimension of the template, obtained by taking the dot product of the calcium response with the stop template. Landmark stops have a higher coefficient than spontaneous stops, indicating that the stop template has a greater contribution to responses during landmark stops. **(G)** Box plots for the distribution of the coefficients in (F). The orange lines represent the respective medians. The means of the distributions are significantly different (Paired t-test, ***:p=1.09e-6). **(H)** Examples of the calcium activity of six M2 neurons projecting to STN during stops at the landmark and in the middle of the track. The response for spontaneous-stops is normalized to the maximum value of the corresponding response for landmark-stops. **(I)** An AAV virus expressing NpHR3.0 was bilaterally injected in M2 of wild-type mice (N=5) and a optic fibers were bilaterally implanted over STN to optogenetically target M2 axons in STN. On pseudorandom trials, amber light (589nm) was presented continuously starting position 175 for either 1.5 seconds or till the animal crossed position 275, whichever occurred first. **(J)** Sagittal section showing M2-STN projections expressing EYFP and NpHR3.0. **(K)** Plot showing the fraction of miss trials during laser ON (inhibiting M2-STN axons), during laser OFF trials (Mann-Whitney U test, *:p=0.045<0.05), and during laser OFF trials for control experiments expressing GFP instead of NpHR3.0 (Mann-Whitney U test, ***:p=0.0<0.005).

For a principled clustering, we derived a low dimensional subspace which explained more than 80% of the energy in the neural population response (**Figures 4B and S3A,B**). Through a change of basis, each neuronal response corresponded to a weighted combination of three basis functions (**Figures 4C and S3A,B**), representing ideal pre-stop, stop and post-stop neurons, with an additional noise term **(Methods)**. We used the weights to cluster the neurons into three groups and recovered a fraction that is active during stops (**Figure 4D**), which showed significantly more reliable responses to stopping compared to the other two groups (**Figure 4E**). We then collected activity during spontaneous stops performed by the animal as it ran towards the landmark. We aligned the activity to switching points in spontaneous-stop windows (as done for landmark-stop windows) and averaged it across windows for stop neurons. We found that M2 neurons are significantly more active at stops at the landmark that are visually-guided compared to spontaneous stops in the middle of the track (Paired t-test, p=1.09e-6) (**Figures 4F-H and S4A-C**), indicating that during task performance M2 neurons specifically signal goal-directed visually-guided stops.

Our imaging results revealed a surge of activity at the onset of visually-guided stop and our axonal activation results showed that such a surge can halt locomotion. To assess whether such an activity surge is necessary, we expressed NpHR bilaterally in M2, and implanted two optic-fibers bilaterally over STN to target the M2 axons there (**Figure 3I, J**). On a random subset of trials, we delivered continuous amber (589nm) light to inhibit the axons when the animal approached the landmark. We found that inactivation increased the number of misses (**Figure 3K**), while light delivery in control experiments (**Figure S2D,E,G,H**) does not. With the role of the surge of activity established, the question we then asked was: how does that surge of activity drive locomotion to halt?

### Behavioral dynamics can be physiologically realized through feedback control

Our behavioral model indicates that the animal picks an ON-OFF locomotion plan that drives the velocity of the animal. Crucial in that locomotion plan is the switching point, upon which the animal initiates fast braking. Our physiological and optogenetics experiments showed neuronal stop signals that may lead the animal to initiate stopping, at the switching point. The question remains as to how these stop signals enable locomotion halts. Particularly, we next sought a physiological realization of the behavioral model that connects the physiology and anatomy to the braking dynamics as depicted by the parameter τ. We modeled the physiological dynamics through a feedback control system, whereby the neural circuitry tracks a neuronal reference signal depicting the locomotion plan and ensures a quick reaction in velocity at the onset of stopping. To define such a control diagram (**Figure 5A,B**), we needed to define the system we are controlling and the controller.

**Figure 5:**
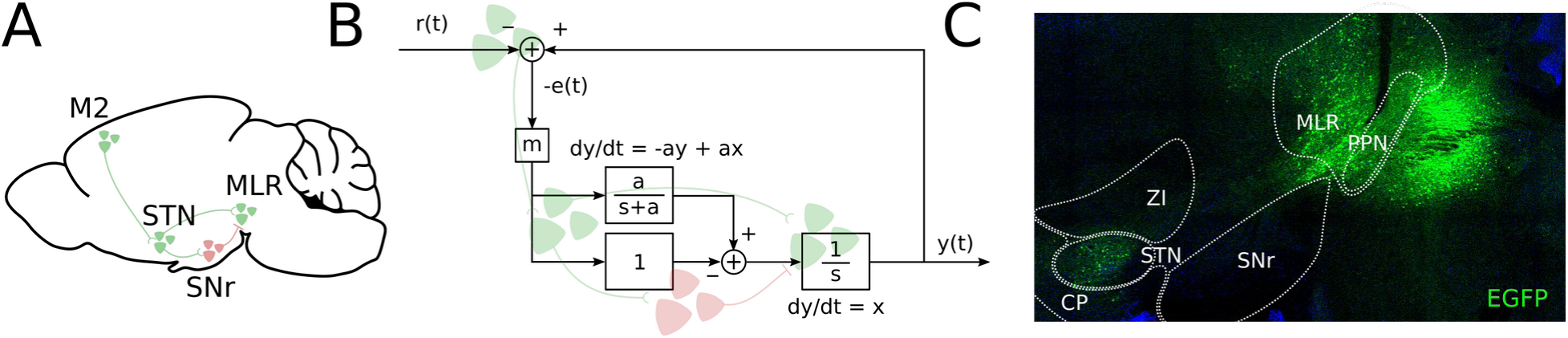
The behavioral dynamics can be physiologically realized through feedback control. **(A)** Schematic showing the implicated neural circuit with green and red indicating excitatory and inhibitory cells, respectively. Activity through the cortico-subthalamic projection can reach the MLR either through a direct STN-MLR projection or via an STN-SNr-MLR pathway. **(B)** Control theoretic model of cortico-subthalamic activity that enables rapid control of locomotion. Each box indicates a transfer function in the Laplace s-domain and is labeled with the corresponding input-output relation with x, y and t denoting input, output and time. The pathways bifurcating from STN interact to simulate mathematical differentiation, cancelling the slow integrative dynamics of the MLR. **(C)** Sagittal slice showing STN vglut2+ neurons projecting to MLR. A AAVrg-EF1a-DO_DIO-tdTomato_EGFP virus was injected in MLR of vglut2-cre animals (N=3) to retrogradely express EGFP in vglut2+ neurons projecting to MLR.

The mesencephalic locomotor region (MLR) in the midbrain is an evolutionarily-preserved structure whose electrical stimulation in cats elicits locomotion at a range of speeds and gaits, scaling with the applied stimulation frequency (Ryczko and Dubuc, 2013; Shik et al., 1969; Whelan, 1996). Such effects have been recapitulated in mice (Roseberry et al., 2016) and optogenetic manipulation of the MLR glutamatergic population has been shown to bidirectionally control locomotion (Caggiano et al., 2018; Josset et al., 2018; Roseberry et al., 2016). We thus considered the glutamatergic population of the MLR as the system we are controlling. Electrical and optogenetic stimulation of the MLR shows that activity and speed is sustained beyond stimulation (Roseberry et al., 2016). The activity decay and locomotion stopping that occurs in such a setting is on the order of seconds after stimulation. However, in our task, locomotion halts occur within the first 100ms of our behavioral switching point. Because of this short temporal period, we can computationally assume that MLR activity does not decay and therefore model the MLR as an ideal integrator, integrating neuronal input to drive the dynamics.

The basal ganglia have been shown to decrease MLR activity via the indirect pathway (Roseberry et al., 2016), notably through SNr (Freeze et al., 2013; Ryczko and Dubuc, 2013; Liu et al., 2020). Information from the STN reaches the MLR through SNr, supporting an inhibitory role of STN onto MLR. Furthermore, STN has direct excitatory projections to MLR (Roseberry et al., 2016; Caggiano et al., 2018), which we additionally verified by retrograde viral tracing (**Figure 5C**). Together, these projections have a crucial role in our model for establishing fast input-output dynamics (**Figure 5A**). The direct STN projections to MLR are understudied and their functional role is unclear (Hamani et al., 2004), and this served as a motivation for our investigation. More generally, SNr projects diffusely to various brain structures, and while we are positing that the SNr pathway to MLR controls locomotion, control of other actions are likely mediated via pathways to structures other than MLR.

Our model proposes M2 as computing a discrepancy between a locomotion plan and the current locomotion state of the animal (**Figure 5B, Text S2**). At the switching point, derived from the behavioral model (**Figure 2A**), the discrepancy is significant, leading to ‘error-signals’ that are sent down along the M2-STN projections. These error signals appeared thus far in the form of stop signals, notably in the imaged responses. Signals of a possibly similar nature have been reported in frontal cortices, to report visuomotor mismatch, in the context of predictive coding (Attinger et al., 2017; Heindorf et al., 2018). Since we defined the MLR as integrating neuronal input to drive locomotion, feeding these error signals directly to MLR would correct its response and allow a decrease in velocity. Crucially, however, if these error signals were directly fed into MLR as a control mechanism, the animal will not be able to stop quickly enough to collect reward. Indeed, the behavioral model has the parameter τ that is required to be small enough for locomotion halts to be rapid. We propose that the controller needs to overcome the slowness limitation by performing a mathematical differentiation operation, thereby cancelling out the slow integrative dynamics of MLR. Anatomically, STN projects to MLR via two pathways, one inhibitory and one excitatory, whose precise temporal interaction can simulate differentiation, yielding the necessary computation to drive rapid dynamics (**Figure 5A,B**). To fully characterize the controller, we require three kinds of information: on the input space, the input-output relation and the dynamical state of the controller. The physiological experiments that follow are designed to provide this information (**Figure S5A-C**).

We purposefully kept the model simple to focus it on a key point: capturing the speed of stopping through a time constant. The main prediction of the model is the necessity of a controller that performs differentiation (essentially a high-pass filter) of the M2 stop signals; without such a controller, we cannot attain an adequate time constant. The next experiments are designed to examine this point.

### Fast input-output dynamics are enabled by mathematical differentiation

The two-photon calcium imaging of STN-projecting M2 neurons provided us with the information needed to characterize the input space. To study the input-output relation, we recorded single-unit extracellular activity in M2 and MLR simultaneously, using silicon probes. Our goal was to recover the neurophysiological signals arising at the input and the output of the controller, and specify through the input-output relation the computational characteristics that are required by a controller to achieve rapid locomotion halts. We transition from 2-photon imaging to single-unit extracellular recordings in M2 and MLR to gain the temporal resolution required to assess input-output dynamics, and analyze the decay in neural responses at the animal stops locomoting. While we lose the cell specificity by doing so, we recover it through additional opto-tagging experiments.

For each recorded unit, we retrieved the spiking activity during the landmark-stop windows, averaged all of the activity, then smoothed it to obtain average firing rates ( **Methods**). Sorting the average responses revealed a heterogeneity in the types of neural responses in both M2 and MLR (**Figure 6A**), similar to that observed by calcium imaging of M2 responses (**Figure 4A**). We then performed a principled clustering of neuronal responses, as performed for the imaged responses (**Figure 4B,C**), and similarly recovered three groups of neurons, revealing a fraction that are active during stops (**Figure 6B,C**). Both brain structures are considered to be implicated in the task and admit such stop neurons; however the fraction of stop neurons is significantly higher in M2 (51/126 in M2 versus 17/141 in MLR; chi-square test p=3.5e-14), indicating that these signals are not equally widespread throughout the brain (**Figure 6C**). Furthermore, each group of neurons was found to respond more reliably within its corresponding epoch, when compared to the other groups (**Figure 6D**).

**Figure 6:**
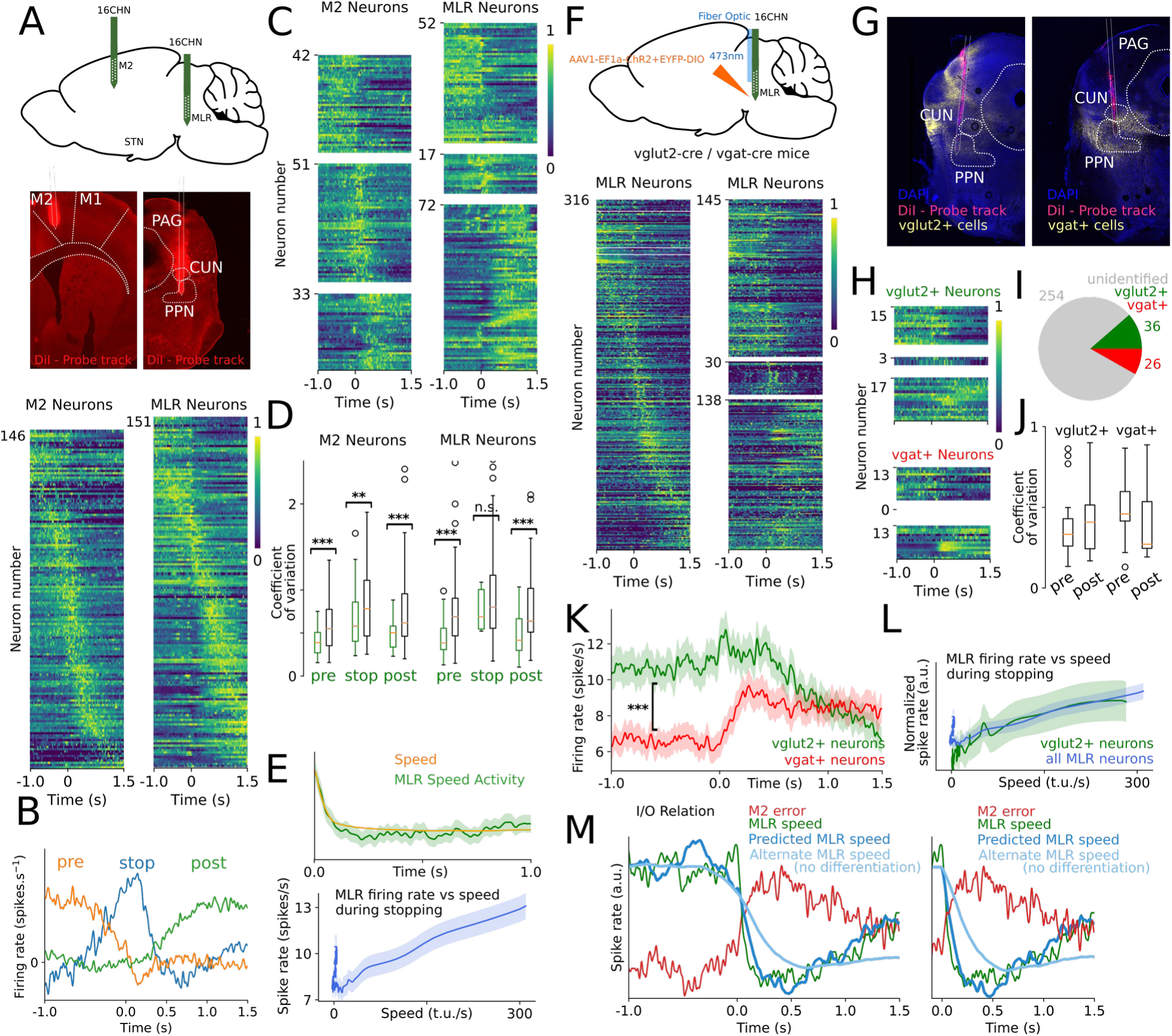
Fast input-output neuronal dynamics are enabled by mathematical differentiation. **(A) Top.** We recorded extracellular single-unit activity using two 16-Channel silicon probes, simultaneously, in M2 and MLR of wild-type mice (N=4). **Middle.** Coronal sections showing DiI recording probe track in M2 and MLR. **Bottom.** Plot showing the normalized average firing rate of M2 and MLR neurons, within landmark-stop windows aligned to the switching point. **(B)** Graph showing 3 templates of ideal neuronal responses related to three epochs in the task: pre-stop, stop and post-stop. Each neuronal response can be expressed as a weighted combination of these 3 templates. **(C)** Plots of all the neurons whose neural response energy (area under the squared signal) was more than 80% explained by the subspace, clustered into three groups (pre-stop, stop and post-stop, from top to bottom) using the templates in (B). The stop-related activity is significantly more prominent in M2 compared to MLR, comprising 40.5% (51 out of 126) of the neurons in M2 versus 12.1% (17 out of 141) in MLR (chi-square test, p=3.5e-14<0.005). **(D)** Boxplots showing the reliability of neuronal responses during the pre-stop, stop and post-stop period for M2 and MLR neurons. The reliability of each of the three clustered population in (C) is computed in its corresponding epoch (e.g., the reliability of the pre-stop cluster is computed at the pre-stop epoch) and compared to that of the remaining neurons. High reliability is measured by a low coefficient of variation. Clustered neurons show increased reliability for their corresponding epoch (Mann-Whitney U test, ***:p<0.005, **:p<0.01), with stop MLR neurons showing a non-significant difference from the remaining neurons (Mann-Whitney U test, p=0.329). **(E) Top.** Plot superposing the speed of the animal and the speed-related MLR neural response following the switching point. **Bottom.** Plot showing the speed-related MLR neural response plotted against the speed of the animal, following the switching point. The dashed red line is fit through linear regression to the data during the first second after the switching point (R^2^=0.829 p=5.36e-78). Activity in MLR is linearly related to the animal’s speed during locomotion halts. **(F) Top.** ChR2 was expressed in a cre-dependent manner in MLR neurons of vglut2-cre and vgat-cre mice. An optic fiber coupled with a recording probe was lowered above MLR to identify vglut2+ and vgat+ MLR cells expressing ChR2 while recording. **Bottom.** Plots showing all the neurons recorded during phototagging and their clustering into three clusters as in (C). **(G)** Coronal sections showing DiI probe track location and ChR2 expression (mCherry) in vglut2+ and vgat+ MLR cells. **(H)** Plots showing the three clusters of vglut2+ and vgat+ identified cells as clustered in (C). **(I)** Plot showing the fraction of unidentified, vglut2+ and vgat+ cells among recorded cells. **(J)** Boxplots showing the reliability of neuronal responses for pre- and post-stop neurons during the pre- and post-stop epochs, respectively. Vglut2+ neurons show higher reliability in pre-stop neurons (Mann-Whitney U test, n.s.:p=0.325), while vgat+ cells show higher reliability in post-stop neurons (Mann-Whitney U test, n.s.:p=0.129). **(K)** Plot showing the average responses of vglut2+ and vgat+ neurons, with a significant difference in the pre-stop epoch (Mann-Whitney U test, ***:p=1.6e-37<0.005). **(L)** Plot showing the normalized speed-related MLR neural response of vglut2+ cells (green) and the cells recorded in (A) (blue) plotted against the speed of the animal, following the switching point. The linear relation observed in (E) extends to vglut2+ neurons. **(M) Left.** Plot showing the reconstructed M2 error signal (input), the MLR response (output) and the predicted MLR response using the reconstructed input-output relation (predicted output) starting from 1s before the stopping onset. The plot also shows an alternative prediction obtained by removing the STN-MLR projection from the model, where we find that activity cannot decay quickly enough (RMSE computed starting from the switching point, RMSE_alternate_/RMSE_predicted_=4.45 fold increase). **Right.** Plot showing the same analysis but performed starting at the onset of stopping, by forcing the error signal to be zero before time 0, thereby forcing speed decay to initiate only at the onset of stopping (RMSE_alternate_/RMSE_predicted_=6.63 fold increase).

Our behavioral model pertains to controlling the velocity of the animal. Our physiological model pertains to controlling a neurophysiological signal reflecting the velocity of the animal. If we are analyzing the decay in velocity through a decay in neuronal activity, then it is necessary to verify that the velocity and the corresponding neural activity do not produce time lags or decay at different rates. To investigate this, we recovered neural activity in MLR that reflects locomotion speed (**Methods**) and compared it to the decay in speed. We observed that the activity is linearly related (up to baseline offset), thereby incurring no change in decay rates (linear regression during the second after onset of stopping, R^2^=0.829 p=5.36e-78) (**Figure 6E**). This implies that we can study the average rate of decay in velocity during stopping, by studying the average rate of decay in neural activity during stopping.

The MLR however has a number of cell-types, with different, potentially opposing, functions (Caggiano et al., 2018; Josset et al., 2018). Thus, we next investigated the roles of excitatory and inhibitory cells in the MLR in the context of the task using optotagging. We expressed ChR2 in a cre-dependent manner in MLR of vglut2-cre and vgat-cre mice to identify excitatory and inhibitory cells, respectively. We lowered a recording probe coupled with an optic fiber, and identified the photo-responsive responsive cells (Figure 6F,G). The recordings and clustering (Figure 6F) revealed similar activity and proportions obtained in the initial recordings (Figure 6A,C). The recordings gathered 36 vglut2+ units and 26 vgat+ units (Figure 6I), and showed responses in both pre-stop and post-stop epochs. The reliability of the responses between populations was different: pre-stop vglut2+ were more reliable in the pre-stop phase than were the post-stop vglut2+ neurons in the post-stop phase (Figure 6H). The reverse was found to be true for vgat+ cells. This supports the findings that excitatory cells promote whereas inhibitory cells suppress locomotion. To further investigate this, we averaged the trial responses and observed a drastic difference: the response of vglut2+ cells decayed following the stop while that of vgat+ cells increased (Figure 6K). The general pattern of spiking activity of vglut2+ cells follows that observed in the single unit recordings shown in Figure 6E, and overlaying the plots (after affine transformation to account for differences in firing rates and baseline activity) showed that the two are closely matched (Figure 6L).

We can derive from our physiological model (**Figure 5A**) the characterization:

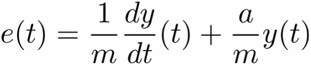

The error signal consists of a weighted combination of acceleration and velocity (**Text S2**). However, the theoretical error signal around the switching point is negative, as *r(t)* becomes *0* while *y(t)* is positive. As neural firing rates are fundamentally non-negative, the error signals sent from M2 to STN can only correspond to *-e(t)*, the negative of the error term. This, in turn, consists of a weighted combination of a negative acceleration signal *-dy/dt* and a negative speed signal *-y* (**Text S2**). We then reconstructed these signal components in M2 (**Methods**) and reconstructed the input-output relation by identifying the coefficient *a* (=17.08). Using the reconstructed input-output relation, we derived the predicted MLR response (**Figure 6M**). Crucially, we constructed an alternative model by removing the differentiation component, leaving only an amplification gain as a means of control. We found that the animal cannot decrease its MLR activity in a timely manner (**Figure 6M**), suggesting the necessity of a controller performing additional processing to speed up the dynamics (RMSE computed starting from the switching point, RMSE_alternate-response_/RMSE_predicted-response_=4.45 fold increase).

### STN supports the dynamical state required to drive the dynamics

Our derived input-output relation suggests that a differentiation operation is implemented prior to the MLR input to ensure fast input-output dynamics. This relation relies on delays, and therefore requires a dynamical state to support it. This state is a one-dimensional variable θ (**Text S2**) governed by:

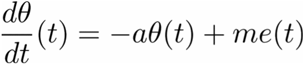

The output of the MLR then corresponds to integrating the difference between the dynamical state and the input to STN:

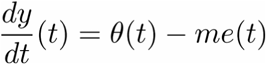

Our physiological model further suggests that differentiation is implemented by sending elements of *θ(t)* and *me(t)* along two complementary pathways from STN to MLR. As the existence of information related to these two signals forms the basis of differentiation, we performed extracellular single-unit recordings in STN to reconstruct elements of the error signals, the dynamical state and the differentiation operation (**Figure 7A, B**).

**Figure 7:**
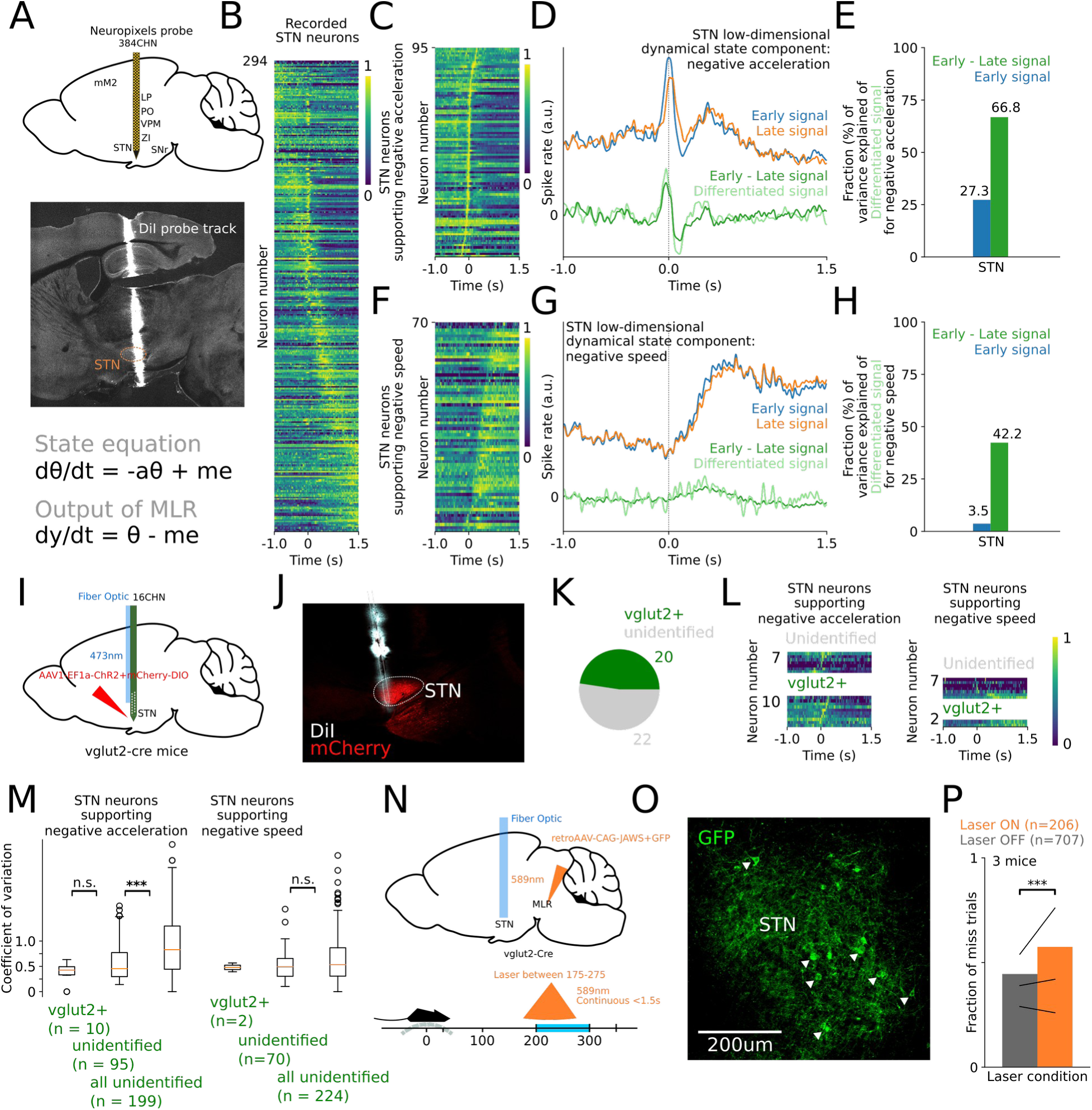
STN supports the dynamical state required to drive the dynamics. **(A) Top.** We recorded extracellular single-unit activity in the STN of wild-type mice (N=4) using Neuropixels probes. **Middle.** Sagittal section showing the Neuropixels probe placement track (DiI) after recording. **Bottom.** Equations that dictate the evolution of the dynamical state *θ*, and its interaction with the error signal *e* to produce a differentiated input *y* to the MLR. The scaling factors *a* and *m* represent the time-constant and gain in the control diagram of Figure 5B. **(B)** Plot of the normalized firing rate of STN neurons, within landmark-stop windows aligned to the switching point. **(C)** Plot showing neurons whose activity peaks between 250ms and after 250ms of the stop onset. The neurons are ordered by peak timing. **(D)** Within the population of (C), we recreated two low-dimensional signals representing the negative acceleration component of the error signal (early signal), and its dynamical state counterpart (late signal). The difference of these two produced a differentiated signal, matching the theoretical prediction. **(E)** Bar graph showing the variance of the theoretical differentiated signal in (D) explained by the difference between the Early and Late signals (green) versus that explained by the Early signal as if no computation was performed (blue). **(F)** Plot showing neurons whose activity transitions from low to high between 250ms before the stop onset and 750ms after. **(G)** Within this population of (F), we recreated two low-dimensional signals representing the negative speed component of the error signal (early signal), and its dynamical state counterpart (late signal). The difference of these two produced a differentiated signal, matching the theoretical prediction. **(H)** Bar graph showing the variance of the theoretical differentiated signal in (G) explained by the difference between the Early and Late signals (green) versus that explained by the Early signal as if no computation was performed (blue). **(I)** ChR2 was expressed in a cre-dependent manner in STN neurons of vglut2-cre mice (N=2). An optic fiber coupled with a recording probe was lowered above STN to identify MLR cells expressing ChR2 while recording. **(J)** Sagittal section showing the DiI probe track location and ChR2 expression (mCherry) in vglut2+ cells in STN. **(K)** Plot showing the fraction of vglut2+ identified cells among recorded cells. **(L)** Plots showing neuronal responses clustered as in (C) and (F) for the recorded cells. **(M)** Boxplots showing the reliability of neuronal responses during the stop period for STN cells supporting negative acceleration, and during the post-stop period for STN cells supporting negative speed. Vglut2+ cells show a reliability similar to the unidentified cells during the stop period (Mann-Whitney U test, n.s.:p=0.236), which show significant difference in reliability compared to the remaining recorded STN cells, not supporting negative acceleration (Mann-Whitney U test, ***:p=5.4e-7<0.005). **(N)** A retroAAV expressing JAWS in a cre-dependent manner was bilaterally injected in MLR of vglut2-cre mice, and optic fibers were bilaterally implanted over STN to target the vglut2+ STN neurons projecting to MLR. On pseudorandom trials, amber light (589nm) was presented continuously starting position 175 for either 1.5 seconds or till the animal crosses position 275, whichever occurs first. **(O)** Sagittal section showing STN neurons projecting to MLR and retrogradely expressing GFP and JAWS. **(P)** Plot showing the fraction of miss trials during laser ON and OFF trials (Mann-Whitney U test, ***:p =0.004 < 0.005).

We studied the dynamical state of the controller by linearly decomposing it into two components, as done for the error signal: one component employed to differentiate the negative acceleration component and another component employed to differentiate the negative speed component (**Text S2**). We first restricted the neuronal population to the neurons in STN whose firing activity peaked between 250ms before and 250ms after the switching point (**Methods**) (**Figure 7C**). We considered 250ms after the switching point as MLR speed activity had reached its minimum at approximately that point (**Figure 6D**), and we considered 250ms before the switching point for symmetry and to capture peaks that occur immediately before braking. We found two dimensions in a low-dimensional neural activity space where the first encodes the negative acceleration, and the other encodes its corresponding dynamical state. Crucially, the difference of the activity along these dimensions yields the needed differentiated signal of negative acceleration (**Figure 7D, E**). We repeated the same procedure for the population of neurons in STN whose firing activity transitioned from low to high after 250ms (**Figure 7F**) (**Methods**), the waiting phase as determined by MLR activity (**Figure 6D**), and again were able to reconstruct two dimensions that yield the differentiated signal of the negative speed component (**Figure 7G, H**). Combining the subspaces for these two populations indicate that activity in STN can support differentiation.

We then asked: is this information a characteristic of STN activity, or does it exist in other brain areas? We thus repeated the process for the M2 extracellular recording data. The best derived two-dimensional subspace could not reconstruct the dynamical state component corresponding to the negative acceleration component in the error signal (66.8% versus 27.2% of the theoretical signal variance explained by STN and M2, respectively) (**Figure S6A-C**). This negative result also held as we increased the initial low-dimensional subspace we searched in by doubling the number of dimensions (**Figure S6A,D**). Indeed, the neural stop-related activity in STN showed a very fine tuning to stopping (**Figure 7C**), unlike that of M2 activity (**Figure S6B**). However, less temporal precision is required to represent the negative speed signal, and we found some correlates in M2 for the dynamical state corresponding to it (42.2% versus 28.2% of the theoretical signal variance explained by STN and M2, respectively) (**Figure S6D-H**). However, it is the negative acceleration component that drives the dynamics to halt initially, and the negative speed component is mainly required to sustain the halt by canceling out rebound effects from the negative acceleration component.

To verify that these stopping dynamics are indeed related to STN neurons, we photo-identified STN cells by expressing ChR2 in STN of vglut2-cre mice and delivering light while recording (Figure 7I,J). Our recordings have identified 20 vglut2+ cells in 2 mice (Figure 7 K), whose activity mostly supports negative acceleration, corresponding to the stop signal (Figure 7L,M). This justified studying the unidentified cells in the stopping regime.

Our experiments then suggest a role for the STN-MLR projections in interacting with STN-SNr-MLR pathway to accelerate the signals. To test this, we bilaterally injected in MLR of vglut2-cre mice a retroAAV that expresses JAWS in a cre-dependent manner (Figure 7N). This causes JAWS to be expressed in STN somas projecting to MLR, among other population. We then bilaterally implanted optic fibers above STN to target these projection neurons (Figure 7O). On a random subset of trials, we delivered amber light when the animal was approaching the landmark, and found that inhibiting these projections led to an increase in miss rate (Figure 7P). This projection is excitatory, and is known to project to excitatory cells in MLR. This the counterintuitive role in accelerating stopping (rather than promoting locomotion) via its interaction with the STN-SNr-MLR pathway.

## Discussion

We have used a simple visually guided locomotion task to derive an important principle of movement control that employs a specific pathway, from M2 to STN, and circuit, from STN to MLR. Our results demonstrate that a signal is sent down along M2-STN projections to rapidly halt locomotion. Importantly, these signals are visually guided and do not occur during spontaneous locomotion halts. Our results further suggest that the M2-STN-MLR pathway ensures this fast response by differentiating the signal, to compensate for integration dynamics in MLR. In particular, STN activity can reach MLR through SNr, but also through direct STN-MLR projections; the temporal interaction of these two pathways simulates differentiation.

Our analysis began by capturing the behavior of the animal through a minimum-time optimal-control problem, which has been extensively studied in the literature to yield a bang-bang optimal control solution (Bertsekas, 1995). This approach captured the rate of stopping through a time-constant, and made the notion of switching point central to the analysis. This approach then enabled us to cast the goal of the neural signaling as ensuring a small enough time constant to achieve rapid locomotion halts. We formalized the neural signaling through a feedback control system, and, through a set of experiments, characterized the properties governing the input space, the input-output relation and the dynamical state of the controller.

Populations of neurons in the brainstem, notably in the medulla, have been found to stop ongoing locomotion (Bouvier et al., 2015; Capelli et al., 2017; Grätsch et al., 2019; Juvin et al., 2016). We expect such reticulo-spinal cells to be recruited in the context of our task. We do not think that the cortico-subthalamic signaling through MLR is bypassing these lower brainstem circuits, and this warrants detailed future investigations. Yet, the nature of the lomotion signaling is different. These reticulo-spinal cells impinge directly onto spinal circuits, and offer a low-level direct control of locomotion. However, the cortico-subthalamic signaling necessitates further processing to reach locomotor circuits, and the processing should be designed so that it ensures fast locomotion halts. From an engineering perspective, feedback control can be leveraged to steer a system’s trajectory as needed and ensured desired operation, which in our case consists of additionally ensuring fast responses. This principle is often realized by computing an error signal—reflecting the discrepancy between a reference trajectory and the current trajectory of the system—and employing it to oppose the deviation of the system from a desired operation (Aström and Murray, 2010). Our behavioral model implicates a locomotion plan as a reference signal, from which such error signals can potentially be derived. Error signals have been reported widely in the brain. They provide, for instance, a basis for the dopaminergic reward system (Schultz, 1998), and have been instrumental in the control of movement (McNamee and Wolpert, 2019; Shenoy et al., 2013; Wolpert and Ghahramani, 2000), setting a basis for optimal feedback control in motor coordination (Todorov, 2004; Todorov and Jordan, 2002). Error signals necessitate reference trajectories, and such references are often derived from predictive capabilities of brain function. These capabilities have often been formalized through internal models (Huang et al., 2018; McNamee and Wolpert, 2019) and predictive processing (Keller and Mrsic-Flogel, 2018). Indeed, error signals have been especially reported following expectation perturbations, notably visuomotor mismatch (Attinger et al., 2017; Heindorf et al., 2018; Marple-Horvat et al., 1993), as signaling prediction errors. The signals we elucidate could also be of a similar nature. The mechanisms by which they arise are the subject of further research.

Stopping in such a task is typically considered to recruit proactive inhibition as opposed to reactive inhibition. Indeed, the animal can see the landmark as it approaches, and can prepare to stop. The situation then lends itself to a potential locomotor plan that is implemented without being interrupted. The role of the hyperdirect pathway has been extensively studied in reactive settings (Aron et al., 2003; Eagle et al., 2008; Nachev et al., 2008, 2007). In stop-signal reaction tasks or go/no-go tasks, participants are signaled to immediately halt an ongoing (or to-be-initiated) action. Proactive inhibition is considered to be heavily mediated by the indirect pathway, but there is certainly evidence of the hyperdirect pathway, and more generally the same ‘stopping network’, as equally being involved (Aron, 2011; Meyer and Bucci, 2016). Our work also highlights a role for the hyperdirect pathway as a critical route for rapid cortical modulation of brainstem structures, complementing the classical role of the direct and indirect pathway, and offering a view consistent with how the hyperdirect pathway is considered to interact with them (Schmidt et al., 2013). We believe that the hyperdirect pathway does indeed have a general role in interrupting action (Aron et al., 2016, Fife et al., 2017, Li et al., 2020). However, aspects of that role need not come through the pathway to MLR, but perhaps through pathways to the superior colliculus (SC) or thalamus through SNr. One can hypothesize that the role of the corticosubthalamic pathway towards SC might be in rapidly interrupting orienting behavior. The role of the hyperdirect pathway may well extend beyond motor, to associative and cognitive processes. Evidence of this generality is emerging (Heston et al., 2020; Hannah and Aron, 2021), and this role is still a subject of further research.

In addition to the indirect pathway in the basal ganglia, alternative routes for signaling locomotion halts might include a visual tectal pathway, consisting of direct projections from the superior colliculus to the mesencephalic locomotor regions (Roseberry et al., 2016). We expect such a signal to be more engaged upon sudden flashes of the landmark, likely in a more reactive setting, perhaps engaging circuits typically used for startle responses (Liang et al., 2015). Additionally, glycinergic neurons in the pontine reticular formation (PRF) project to the intralaminar thalamic nuclei, and stimulation of their axons in IL produces behavioral arrest (Giber et al., 2015). M2 directly projects to PRF, and the projections might directly drive such glycinergic cells to achieve fast locomotion halts. Aside from these pathways, various other pathways can induce behavioral arrest (Klemm, 2001; Roseberry and Kreitzer, 2017). These are generally recruited via different mechanisms, though some can overlap with ours. Further task enhancements are needed to elucidate their contributions.

## Acknowledgment

This work was supported by NIH grants NS090473, EY007023 and EY028209 (M.S.) and a JPB Foundation Fellowship (E.A.). We thank Dr. Vincent Breton-Provencher and Dr. Grayson Sipe for their careful reading of the manuscript, and members of the laboratory for their comments and advice.

## Author Contributions

E.A. and M.S. conceived the project and developed the concepts presented. E.A. designed the experiments and performed them with assistance from T.J. in behavioral training. E.A. analyzed the data and developed the theory. E.A. and M.S. wrote the manuscript with input from T.J..

## Methods

### Animals +

All procedures were approved by the Massachusetts Institute of Technology’s Animal Care and Use Committee and conformed to the National Institutes of Health guidelines. Adult mice (>2 months old) on a C57BL/6J background were used in this study. Male or female mice were randomly selected for each experiment. We also used the Ai148D (Ai148(TIT2L-GC6f-ICL-tTA2)-D, Jackson Laboratory) mouse line for the two-photon imaging experiments, the vglut2-Cre (Vglut2-ires-cre knock-in, Jackson Laboratory) mouse line for the optogenetics, optotagging experiments and tracing experiments and the vgat-Cre (Vglut2-ires-cre knock-in, Jackson Laboratory) mouse line for the optotagging experiments.

### Surgery

Surgical procedures were performed under isoflurane anesthesia while maintaining body temperature at 37.5 °C using an animal temperature controller (ATC2000, World Precision Instruments). After deep anesthesia was confirmed, mice were placed in a stereotaxic frame (51725D, Stoelting), scalp hairs were removed with hair-remover cream, the underlying skin was cleaned with 70% alcohol and betadine, and an incision was made in the scalp. The conjunctive tissue was removed by rubbing hydrogen peroxide on the skull. The skull was positioned such that the lambda and bregma marks were aligned on the anteroposterior and dorsoventral axes. The skull was further leveled along the mediolateral axis. Animals were given analgesia (slow release Buprenex, 0.1mg/kg and Meloxicam 0.1mg/kg) before and after surgery and their recovery was monitored daily for 72 hours.

For viral injections, a 200-μm diameter hole was drilled through the skull at the location of interest. Viruses were delivered with a thin glass pipette at a rate of 75 nL/min (unless indicated otherwise) by an infuser system (QSI 53311, Stoelting). The following viruses (titer: ∼10e-12 virus genomes per ml) were injected in the performed experiments: AAV5-CamKII-ChR2-mCherry (pAAV-CamKIIa-hChR2-(H134R)-mCherry-WPRE-pA, UNC Vector Core) for optogenetics activation of M2 axons in STN, AAVrg-EF1a-Cre-mCherry (pAAV-Ef1a-mCherry-IRES-Cre, Addgene) for 2P imaging of M2 neurons projecting to STN, AAV5-CaMKII-NpHR3.0+EYFP (pAAV-CaMKIIa-eNpHR 3.0-EYFP, Addgene) for optogenetics inhibition of M2 axons in STN, AAVrg-CAG-Jaws+GFP-FLEX (pAAV-CAG-FLEX-rc [Jaws-KGC-GFP-ER2], Addgene) for optogenetics inhibition of STN somas projecting to MLR, AAV1-EF1a-ChR2+EYFP-DIO (AAV-EF1a-double floxed-hChR2(H134R)-EYFP-WPRE-HGHpA) for optotagging in MLR, AAV1-EF1a-ChR2+mCherry-DIO (pAAV-EF1a-double floxed-hChR2(H134R)-mCherry-WPRE-HGHpA) for optotagging in STN, AAVrg-EF1a-DO_DIO-tdTomato_EGFP (pAAV-Ef1a-DO_DIO-TdTomato_EGFP-WPRE-pA, Addgene) for anatomical tracing of STN projections to MLR, AAV5-CAG-GFP (pAAV-CAG-GFP, Addgene) for optogenetics control experiments.

For the optogenetic activation experiments, wild-type mice were unilaterally (left hemisphere) injected with 400nL of AAV5-CaMKII-ChR2-mCherry in M2 (centered at AP: +1mm, ML: +0.5mm, DV: +0.5mm). After injection, the skin was sutured and we let mice recover for 4– 6 weeks for optimal opsin expression. In a second surgical procedure, a fiber-optic cannula (Thorlabs) with 200um diameter (0.39NA) was implanted above STN (AP: -1.9mm, ML: +1.5mm, DV: +4.3mm). The optic fiber cannula was held by a stereotaxic manipulator, inserted and attached to the skull with dental cement (C&B Metabond, Parkell). To avoid light reflection and absorption, dental cement was mixed with black ink pigment (Black Iron Oxide 18727, Schmincke). A custom-designed headplate (eMachineShop) was implanted at the end of the surgery for head fixation. For the optogenetics experiment inhibiting the M2 axons in STN, wild-type mice were bilaterally injected with 400nL of AAV5-CaMKII-NpHR3.0-EYFP in M2, and a dual optic fiber cannula (Doric lens) was implanted with the two 200um diameter (0.39NA) fibers bilaterally lowered above STN (AP: -1.9mm, ML: +1.5mm, DV: +4.3mm). For the optogenetic experiment inhibiting the STN projections to MLR, vglut2+ mice were bilaterally injected with 300nL of AAVrg-CAG-Jaws+GFP-FLEX in MLR (AP: -4.7mm, ML: +1.25mm, depth: +3.15mm from pial surface), and a dual optic fiber cannula (Doric lens) was implanted with the two 200um diameter (0.39NA) fibers bilaterally lowered above STN (AP: -1.9mm, ML: +1.5mm, DV: +4.3mm).

For the retrograde tracing of STN projections to MLR, vglut2+ mice were bilaterally injected with 300nL of AAVrg-EF1a-DO_DIO-tdTomato_EGFP in MLR (AP: -4.7mm, ML: +1.25mm, depth: +3.15mm from pial surface). After injection, the skin was sutured and we let mice recover for 4 weeks for optimal opsin expression, before harvesting the brain for histology.

For imaging experiments, Ai148D mice were bilaterally injected with 200-300nL of AAVrg-EF1a-Cre-mCherry in STN (AP: -1.9mm, ML: +1.5mm, DV: +4.6mm) and a cranial window centered was implanted above M2 in both hemispheres. The virus was delivered at a slower rate of 50nL/min slowly to limit the spread as much as possible, to avoid contaminating adjascent brain regions. The location of STN also limits contamination: it is encapsulated by the cerebral peduncle from the anterior side, the ventral side, lateral side and half of the posterior side. We drilled a 3-mm circular window centered over the midline (AP: 1mm) to expose M2 on each hemisphere. We stacked two 3-mm coverslips centered on a 5-mm coverslip (CS-5R and CS-3R, Warner Instruments) and glued the three together with ultraviolet adhesive (NOA 61 UV adhesive, Norland Products). The fabricated window was positioned over the craniotomy and attached to the skull using dental cement (C&B Metabond, Parkell). The dental cement was mixed with black ink pigment (Black Iron Oxide 18727, Schmincke) to block light leakage during imaging. We then attached a custom-designed headplate to the skull for head fixation.

Extracellular single-unit recordings were performed acutely in head-fixed behaving mice. We attached a custom-designed headplate to the skull for head fixation, using a custom-designed stereotaxic arm to align the head plate parallel to the median and dorsal line of the skull during implantation. The head plate was attached to the skull using dental cement. The exposed skull was protected using rapid-curing silicone elastomer (Kwik-Cast, WPI) topped with a fine layer of dental cement. One to two days before recordings, mice were anesthetized with isoflurane and the dental cement and silicone elastomer on the skull were removed. The mouse was placed on the stereotaxic frame and 200-μm diameter craniotomies were performed on top of the recording sites of interest, M2 and MLR, centered around the coordinates of interest. The craniotomy was protected with saline and a piece of Gelfoam (Pfizer). The skull was covered again with silicone and the animal was allowed to recover for at least a day. Viral injection were additionally performed before headplate attachment for optotagging experiments: 300nL of AAV1-EF1a-ChR2+EYFP-DIO were bilaterally injected in MLR of vlgut2-cre and vgat-cre mice (AP: -4.7mm, ML: +1.25mm, depth: +3.15mm from pial surface) for optotagging in MLR and 400nL of AAV1-EF1a-ChR2+EYFP-DIO were bilaterally injected in STN of vglut2+ mice for optotagging in STN (AP: -1.9mm, ML: +1.5mm, DV: +4.7mm).

### Behavioral task, equipment and training

Mice were head-fixed using optical hardware (Thorlabs) and positioned on a custom-fabricated rubber treadmill (LEGO Technic + M-D Building Products) coupled with a rotary encoder (Signswise). The virtual runway was constructed using two parallel rails (Thorlabs) each equipped with high-density LED PCB bars (DotStar, Adafruit). Reward consisted of 5-10ul of water and was delivered through a lick spout using a solenoid valve (Parker). Licks were collected, when needed, using a capacitive touch sensor (Adafruit). The behavioral apparatus was controlled through custom-written code deployed on a microcontroller (Arduino). The microcontroller interfaced, through serial connection, with custom-written scripts (Python) running on a custom-designed multi-purpose computer (operating under GNU/Linux) to execute the task.

The landmark consisted of a contiguous set of LEDs that were lit blue on both rails. The length of the landmark amounted to 6cm, and the position of the animal in track units (t.u.) was referenced to the mouse’s nose. Specifically, the landmark interval 200-300 corresponded to the mouse’s nose being within the lit LED range. Track units (t.u.) were calibrated to have 200t.u. correspond to 12cm. The rotation of the treadmill was coupled to the movement of the landmark so that the linear velocity of the outer-edge of the treadmill equaled the linear velocity of the landmark.

Mice underwent water-regulation and obtained water reward (5μl) during behavior. They were then habituated to the treadmill, and underwent a shaping procedure that rewards a stop after crossing a certain distance. The landmark was then introduced and the required waiting time to collect reward was gradually increased across days from 0.6s to 1.5s until they reached their full performance. Experiments began once the number of successful stops at the landmark was consistently above 100 within a 30 minute session.

Our task was designed to elicit short bouts of runs and stops, instead of lengthy periods of locomotion. The aim was to develop symmetry between periods of runs and stops, keeping trial durations short, and to have the time to collect reward depend more on the stopping dynamics. As such, the landmark was positioned relatively close to the animal (15cm) and the length of a trial was then a few seconds (**Figures 1A and S1A**). This design was intended to elicit locomotion in the ‘moderate velocity’ regime (as opposed to high velocity behaviors such as escape behavior) with the goal of engaging the pedunculopontine nucleus (PPN) in the MLR, and more likely basal ganglia circuitry, instead of the cuneiform nucleus (Caggiano et al., 2018; Josset et al., 2018).

The behavioral experiment (**Figure S1J**) assessing the effect of landmark distance was performed by alternating between sessions having the landmark centered at position 250 (near) and sessions having the landmark centered at position 450 (far).

### Behavioral Model

From each behavioral session, we defined landmark-stop windows (rewarded stops between positions 200-300). Each time window corresponds to 2.5s, aligned to a switching point. The switching point was defined as the last major velocity peak (value higher than 25% of the maximum velocity within 200ms window before sustained zero velocity). The time window then consisted of 1s before the switching point, and 1.5s after. We chose this width to capture most of the 1.5s waiting period of the animal, and have the period where speed is decaying to a halt centered, thereby requiring the switching point to be around 1s after the start of the window. Aligning to a switching point, instead of reward-time, allowed us to consider and analyze spontaneous stops that are non-rewarded. We pooled together all landmarks-stop windows, across sessions and animals, and averaged them to get the average traces, presented in (**Figure 2-C**). We then fixed u_min_ = 0 to approximate the problem, and then derived τ and u_max_ from the peak velocity of the animal before stopping and the slope of the logarithm of the velocity trace during decay.

### Optogenetic manipulations

Blue light (473nm) or amber light (589nm) was delivered using a diode-pumped solid-state laser (Optoengine for 473nm, Laserglow Technologies for 589nm). Laser stimulation was triggered using a custom-designed source-follower circuit driven by a microcontroller (Arduino) dictating the stimulation pattern. A fiber-optic patch cable with a ferrule end (200um, 0.39NA) was coupled to the implanted fiber optic cannula with a ferrule mating sleeve (Thorlabs). A piece of black electrical tape was wrapped around the connection between the patch cable and the implanted ferrule, to block any light emitted from that interface.

Photo-stimulation was randomly applied on 30% of the trials. We further imposed a condition that no two consecutive trials could be selected for laser stimulation. For the optogenetics activation experiments, on a trial selected for photo-stimulation, blue light was delivered once the animal reached position 100, for 500ms at 20Hz, 20% duty-cycle (PW:10ms, and T:50ms) with a peak power of 10-15mw (average power of 2-2.5mw). For the optogenetics inhibition experiments, on a trial selected for photo-stimulation, amber light was delivered once the animal reached position 175 continuously for either 1.5 seconds or till the animal crosses position 275, whichever time period is shorter. Each animal underwent three 30min behavioral sessions of photo-stimulation. We kept the session where the behavioral performance was deemed adequate (> 50 hits).

### Calcium imaging and neuronal response analysis

Two-photon calcium imaging was performed through a cranial window. GCaMP6f fluorescence was imaged through a 25x/1.05NA objective (Olympus) using a custom-configured two-photon microscope (MOM, Sutter Systems). Excitation light at 910nm was delivered with a Ti:Sapphire laser (Mai-Tai eHP, Spectra-Physics) equipped with dispersion compensation (DeepSee, Spectra-Physics). Emitted light was bandpass filtered and collected with a GaAsP photomultiplier tube (Hamamatsu). STN-projecting neurons in M2 were imaged between 400-500um below the surface at 10Hz using galvo scanning, and images were acquired by ScanImage (Vidrio) to generate a TIFF stack. Power at the objective ranged from 15 to 30 mW, depending on GCaMP6f expression level and depth.

Neuronal ROI selection and calcium signal extraction was performed using CaImAn (Flatiron Institute) (Giovannucci et al., 2019) implementing a constrained nonnegative matrix factorization approach (Pnevmatikakis et al., 2016). The obtained ROIs and signals were additionally hand-curated to leave out any false-positives.

The animals performed 30min behavioral sessions, and fluorescence was acquired for 1600s (∼26.6 mins) during that period. The behavioral data included a reference signal derived from the microscope acquisition trigger signal, and that signal was used to align to the behavioral signals to the neural signals. The neural signals were then upsampled using piecewise-constant interpolation to match the temporal resolution of 200Hz of the behavior.

#### Data Analysis

From each behavioral session we defined landmark-stop windows (rewarded, stops between positions 200-300) and spontaneous-stop windows (non-rewarded, stops before position 150). Each time window corresponds to 2.5s, aligned to a switching point. The switching point was defined as the last major velocity peak (above 25% of the maximum velocity within 200ms window before sustained zero velocity). The time window then consisted of 1s before the switching point, and 1.5s after.

For each neuron, we computed the average DFF response over a landmark-stop window, normalized each to a maximum DFF of 1, and formed an N-by-T (N: number of neurons and T: time) matrix M where each row corresponded to the normalized average DFF of a neuron. To capture most of the energy in the responses in a low-dimensional space, we reduced neuronal dimensions by performing a low-rank approximation (rank=4). Specifically, we performed a singular value decomposition of M as M=USV^T^ where the matrix S is a rectangular diagonal matrix of singular values, and the matrices U and V are orthonormal matrices (Dahleh et al., 2004; Horn and Johnson, 2012; Strang, 2016), and kept only the highest 4 singular values in S, and set the remaining ones to zero, to get a matrix S_approx_. We reasoned that we had (not necessarily independent) four-degrees of freedom in our analysis: baseline, pre-stop, stop and post-stop activity, and thus sought to begin with a 4-dimensional space. Each of the four non-zero singular values in S corresponded to a temporal neuronal response in V^T^, which together span a 4-dimensional subspace. The subspace contained more than 80% of the calcium signal energy for the whole population (squared Frobenius norm of M). Specifically, the sum of squares of all the entries (energy) in M - M_approx_ is less than 20% of the sum of squares of all the entries (energy) in M, indicating that the low-dimensional subspace indeed captured 80% of the energy in the neuronal responses (Horn and Johnson, 2012; Strang, 2016). Adding more dimensions will only facilitate capturing more of the dynamics by capturing more energy. We aimed to capture the bulk features of the signal, avoiding less prominent features, more fit to be considered noise in these populations, by keeping the number of dimensions to a minimum.

The four temporal neuronal responses obtained in V^T^ corresponding to the non-zero singular values in S_approx_ span a 4-dimensional space, and form a basis to that space. Each DFF response can then be written as a weighted combination of these four basis responses and an additional response outside (orthogonal) to that space that is considered as noise. As a space can admit multiple bases, we decided to find a basis that represents stopping epochs and decompose our neuronal responses onto it. We sought to define ideal neuronal templates of pre-stop, stop and post-stop neurons (**Supplementary Figure 3A**). We began with three signals equal to 1 for t<0 while 0 otherwise, equal to the average velocity of the animal for t>0 while 0 otherwise, and equal to 1 for t>0.5s while 0 otherwise. We projected each of the three square signals to the 4-dimensional subspace defined by the low-rank approximation, and ensured orthogonality using the gram-schmidt orthonormalization process (Strang, 2016). We were left with three templates of ideal pre-stop, stop and post-stop neurons, spanning a three-dimensional subspace. We then kept the imaged neurons for which at least 85% of their calcium signal energy (area of the squared signal) came from the three-dimensional subspace, and discarded the rest from the analysis. This kept about 64% of all neurons we started with, and we performed clustering on them as described below.

Each neuronal response can be written as a weighted combination of the three templates, plus some additional component orthogonal to the subspace (noise). We recovered the weights for each response by taking the inner-product (dot-product) with each template. We multiplied the weight by the maximum value in the template to account for the template width, to get a corrected weight. We then attributed a neuron to one of three classes (pre-stop, stop and post-stop) whose corresponding corrected-weight is highest.

Most importantly, the calcium responses in each class were not reduced in dimensions; they were the original averages of the raw DFF traces taken over landmark stop windows. The dimensionality reduction was only used to cluster the neuronal responses.

For the class of stop-neurons, we computed the weight with respect to the stop-neuron template for the average response in landmark-stop windows and spontaneous-stop windows, by taking the inner-product (dot-product) with the stop-neuron template.

For a fixed epoch and neuron, the activity of a trial was averaged over the epoch and normalized by the average over the whole stopping time window (between -1s and 1.5s, with 0 indicating the switching point). The reliability of a response was the computed as the coefficient of variation (mean / standard deviation). The epochs considered were: pre-stop (-1s to 0s), stop (-0.25s to 0.25s) and post-stop (0.5s to 1.5s).

### Extracellular recordings in M2 and MLR and neural response analysis

On the day of the recording, the animal was head-fixed on the behavioral setup and the silicone and Gelfoam removed gently. A 0.9% NaCl solution was used to keep the surface of the brain wet for the duration of the recordings. We submerged a reference silver wire in the NaCl solution on the skull surface. The 16-channel silicon probes (A1x16-Poly2-5mm-50s-177-A16, Neuronexus) were then lowered in the ventral axis with motorized manipulators (MP-285, Sutter Instrument Company and Micropositioner Model 660, Kopf) at a rate of 20 microns per second. The probe recording in M2 was lowered to (AP: +1mm, ML: +0.5mm, depth: +0.5mm from pial surface) and the probe recording in MLR was lowered to (AP: -4.7mm, ML: +1.25mm, depth: +3.15mm from pial surface). The recording sites on the probe span about 0.375mm. As such, the probe tip was lowered deeper than the indicated depth, to ensure the sites cover the area surrounding the indicated depth.

The extracellular signal was acquired through a custom-configured Plexon MAP system, initially amplified using a 1× gain headstage (model E2a, Plexon) connected to a 100× preamp (PBX-247, Plexon). The signal was high-pass filtered at 300 Hz. Spikes were monitored online with amplitude threshold using Plexon Sort Client software, and raw continuous data was recorded at a rate of 40kHz. Spike sorting was then performed on the raw waveform using a fully-automated spike-sorting algorithm through Mountainsort (Chung et al., 2017). Spike curation was done manually to remove artifacts picked by the algorithms (ill-shaped spikes) and spikes with low amplitudes or low spontaneous spike rate (<0.1 spikes s−1). We verified spike times with cross-correlograms to eliminate duplicates.

For verifying the probe location after recordings, the silicone probes were gently retracted and the recording tract was marked by re-entering the DiI-coated probe (2 mg/ml; D3911, ThermoFisher) at the same location. The brain was harvested post-experiment and the probe location was confirmed through histology.

#### Data Analysis

The animals performed 30min behavioral sessions, and recordings were initiated before the start of a session and lasted after the end of a session. Both the behavioral and physiology apparatus were continuously recording an external reference signal (acquisition at 200Hz in behavioral data and 40kHz in physiology data). This signal was used to align the physiology and behavioral data.

We down-sampled the resolution of the physiology data by binning the spikes into 5ms intervals to match the resolution of the behavioral data, and computed mean average responses as performed for the calcium imaging. The data was smoothed with a Gaussian filter with standard deviation around 7ms (corresponding to a variance of 50ms^2^), and we then performed dimensionality reduction through low-rank approximation (with k=4), exactly as performed for the calcium imaging data.

To study the relation between MLR activity and speed, we considered all the MLR neurons whose average activity transitions from high to low within a 1s window, 750ms before the switching point and 250ms after. Specifically, for each time point, we computed the mean *h* of the activity prior to it and the mean *l* of the activity after it, and found the time point that maximizes *h-l*. We kept the neurons whose optimal time point falls within the corresponding window. We went through all the kept neurons and all stops, paired the spiking activity of one neuron (binned into 5ms intervals to match the behavioral resolution) during a single stop with the position evolution corresponding to that stop window. We then averaged all the position traces, differentiated and smoothed the signal to obtain the speed trace. We similarly averaged all the spiking activity (corresponding to the different stop windows, pooling all the neurons together) and smoothed it to obtain the MLR trace. The smoothing is performed with a Gaussian convolution with standard deviation of about 7ms (var = 50ms^2^). We computed the standard error of the mean error values, similarly. We could then learn the variable *a* by computing the slope of the logarithm of the traces to get a rate of decay of 17.08, approximated thereafter by 17.

To reconstruct the M2 error, we first reconstructed its components, corresponding to the negative acceleration and the negative speed. To reconstruct the negative acceleration signal, we derived the negative acceleration from the speed, and projected it onto the low-dimensional subspace already derived from the neural responses to get the best neural approximation of it. This approximation is achieved by a weighted combination of neural responses where the weights can be either positive or negative. To ensure that the weights are non-negative, we recomputed a new set of weights on the average responses through non-negative least squares, so that a combination of the neural responses using the new non-negative weights matched the low-dimensional response. This yields a low-dimensional representation of the negative acceleration component with only non-negative weights. We repeated the procedure to derive a low-dimensional negative speed signal, beginning by projecting the negative of the speed signal onto the low dimensional space. Finally, the error signal is a positive weighted combination of these two components. To recover these weights, we defined the theoretical impulse response function *ae^−at^u(t)* . We then convolved the separate components with it to get their contribution to the MLR response. Using non-negative least squares, we found the optimal non-negative weights such that the weighted combination of the filtered components best fits the MLR response, leading to the predicted response. By linearity, these correspond to the weights in the original M2 error signal. We then repeated the whole process by removing the STN-MLR projection, and computed a new predicted response via an updated M2 error signal.

The reliability analysis is similar to that performed for the calcium imaging data.

### Extracellular recordings in STN and neural response analysis

The preparation for the STN recordings followed the same procedure as that of M2 and MLR recordings. A motorized manipulator (Micropositioner Model 660, Kopf) was used to lower a Neuropixels 1.0 probe, at a rate of 20 microns per second, towards STN (AP: -1.9mm, ML: +1.5mm, DV: +4.5mm) to a depth of 5.5mm. The spikes analyzed were then restricted to be the ones found on channels corresponding to STN location, taken to be between 4.25mm to 5mm from the pial surface. The data was acquired using the standard Neuropixels hardware (IMEC) though SpikeGLX (HHMI/Janelia Research Campus). Spike sorting was performed using Kilosort2 (Pachitariu et al., 2016), and then hand-curated using Phy.

Probe location was also verified by using DiI after a recording session and harvesting the brain postexperiment to recover the probe track.

#### Data Analysis

As in the M2 and MLR recordings, the animals performed 30min behavioral sessions, and recordings were initiated before the start of a session and lasted after the end of a session. Both the behavioral and physiology apparatus were simultaneously continuously recording an external reference signal (acquisition of 200Hz in behavioral data and 30kHz in physiology data). This signal was used to align the physiology and behavioral data.

The data was again down-sampled by binning the spikes into 5ms intervals to match the resolution of the behavioral data, and was processed as done for the M2 and MLR recordings.

To characterize the component of the dynamical state corresponding to the negative acceleration, we considered all the neurons whose average firing rate is its maximum in a 500ms interval, 250ms before the switching point and 250ms after. We performed a low-rank approximation (k=5) and derived a low-dimensional subspace that explains most of the energy in the average responses. We computed the negative acceleration from the speed signal, and projected it onto this subspace to recover the neuronal signal representing it. We then took this neural signal, and convolved it with *h* = *e^−at^u(t)* to obtain its dynamical state counterpart. This convolution necessarily places the obtained signal outside the low-dimensional subspace, and we then projected it again onto the subspace to ensure that it belongs there. The neural signals representing the negative acceleration component and the dynamical counterpart are both obtained using a weighted combination of the average neural responses via positive and negative weights. To ensure the weights are non-negative, we recomputed a new set of weights (as done in the M2 and MLR data) on the average responses through non-negative least squares, as a means to recover a signal using only non-negative weights that match the low-dimensional response of the non-negative acceleration component and its dynamical counterpart. These non-negative weights span a two-dimensional space, that explained a high amount of energy (39.63%) in the neuronal population response, as it is very close to being a subspace of the original 5-dimensional subspace. We repeated the same procedure to characterize the component of the dynamical state corresponding to the negative speed. For this, we considered all the STN neurons whose average activity transitioned from low to high starting within a 1s interval, 250ms before the switching point and 750ms after the switching point. This was performed using a procedure similar to that used to characterize MLR speed activity.

The reliability analysis is similar to that performed for the calcium imaging data.

### Optotagging in MLR and STN

The preparation for the phototagging recordings followed the same procedure as that of the M2 and MLR recordings. A16-channel silicone optrode (A1x16-Poly2-5mm-50s-177-OA16LP, Neuronexus) was connected via a 105-μm/0.22 numerical aperture patch cable (M61L01, Thorlabs) to a solid-state blue laser (Opto Engine), and was lowered to the recording site of interest as done for MLR and STN. The tip of the optrode was first lowered to around 200um above the recording coordinates of interest, blue light was then periodically delivered (at a rate of 2Hz) while the optrode was then progessively lowered slowly in search for units responsive to light stimulation, adjusting the light stimulation intensity to reduce stimulation artifacts and observe neuronal responses. The final position of the probe was decided when units responsive to light stimulation were detected and the electrode recording sites spanned the recording area of interest.

At the beginning and end of each recording sessions, light pulses of 5 ms at a fixed light intensities (tuned between 0.1-1 mW for a given recording session as a function of evoked responses) were repeatedly delivered in the tissue (frequency: 2 Hz), to perform post-hoc comparison of spontaneous and light-evoked waveform for each sorted unit. Sorting and curation was performed as done for the M2 and MLR recordings. Units were considered light-responsive if they responded significantly using the SALT algorithm (Kvitsiani et al., 2013). We also only kept units responding within an 8-ms-period after light stimulus onset, and whose light-evoked waveforms closely matched the spontaneous ones. Analysis of neuronal responses was performed as done in the respective sections on MLR and STN recordings. The recording tract was also marked by coating the probe with DiI as performed for M2, MLR and STN recordings.

### Histological Methods and Verification

Under very deep anesthesia, mice were perfused transcardially with 0.9% NaCl followed by 4%PFA. The brains were harvested and post-fixed in 4% PFA at 4°C overnight. In some experiments, brains were extracted without transcardial perfusion and only immersed in PFA overnight. Coronal or sagittal sections (100um thick) were cut using a vibratome (VT2008, Leica). Slices were mounted and imaged with a confocal system (TCS SP8, Leica) with 10x/0.40NA or 20X/0.75NA objectives (Leica).

#### Optogenetics

The position and depth of the fiber optic was assessed by delineating tissue damage along the fiber-optic track track following its removal during brain extraction. Placement was considered correct if the M2 axons in STN were found in proximity and within a narrow cone region to the optic fiber tip (0.39NA). Viral expression was considered adequate if cortical projection towards the subthalamic nucleus (through the cerebral peduncle) were strongly visible under the confocal microscope, and fluorescent axons were observed in STN. This applied to activation, inhibition and control experiments regarding the M2 projections to STN. For experiments silencing STN projections to MLR, viral expression was considered adequate if somas were observed in STN and the injection site corresponded to the MLR area, and particularly the PPN per the reference atlas. These verifications were performed using sagittal brain sections.

#### Imaging

The imaging location was determined by identify the center of window, obtained by adequate X- and Y-translations corresponding to the radius of the window starting from an edge, guided by the sinus in the middle of the window, and defining that center as an imaging reference. Imaging fields of view were determined and centered to be around 600um along the ML direction from the origin reference. All imaged neurons in the field of view were considered. This number was further reduced to those whose activity is 85% explained by the low dimensional activity space, as expounded in the section on data processing. Viral expression in M2 was assessed through the 2-photon microscope, and imaging was perfomed when expression showed a dynamic range in fluorescence due to calcium activity. Injection sites were inspected under the confocal microscope through sagittal brain sections, and considered adequated when targetted to the STN, observable under the confocal and additionally visually demarcated by structures observed under the confocal, such that the cerebral peduncle and the SNr.

#### M2 and MLR recordings

The probe recording in M2 was lowered to (AP: +1mm, ML: +0.5mm, depth: +0.5mm from pial surface) and the probe recording in MLR was lowered to (AP: -4.7mm, ML: +1.25mm, depth: +3.15mm from pial surface), which corresponded to the PPN in MLR. The position of the probes was verified under the confocal microscope using DiI, and that of MLR was deemed correct by assessing proximity to the PAG and overlaying a referece atlas to demarcate PPN. These verifications were performed using coronal brain sections. The recording sites of the probe spanned 375um, and as such all sorted recorded units (with more than 500 spikes per session) were included at the start of the analysis. This number was further reduced to those whose activity is 85% explained by the low dimensional activity space, as expounded in the section on data processing.

#### STN recordings

All our neuropixels recordings in STN were performed with the same probe angle, starting from the same coordinates, going to the same depth of 5500um and thereby always reaching the same end-point. These coordinates and angle were chosen during experimental design, using a reference atlas. This defined a fixed recording orientation and direction that crossed multiple brain region, then through the subthalamic nucleus and ended with the tip in the cerebral peduncle. As a check, we indeed observed changes along the probe length once channels were out of the cerebral peduncle, which corresponded to about 500um above the tip. We then recreated this direction on the reference brain atlas, recovered the coordinates of recorded regions per the atlas including the STN, and mapped them to channels on the probe based on the geometry. All sorted recorded units (with more than 500 spikes per session) that we principally detected on recording sites corresponding the STN location were included in the analysis. The position of the probe was verified under the confocal microscope using DiI, and it was deemed correct if it corresponded to the same starting and end point considered by the reference atlas, determined by using brain structures visible under the confocal such as the cerebral peduncle, the STN and the hippocampus. These verifications were performed using sagittal brain sections.

#### Optotagging

In addition to the verifications performed for probe location during extracellular recordings, viral expression was assessed by assessing fluorescence in the recorded region. Verifications for MLR recordings were performed using coronal sections, and for STN recordings using sagittal sections.

## Supplementary Material

**Supplementary Figure 1:**
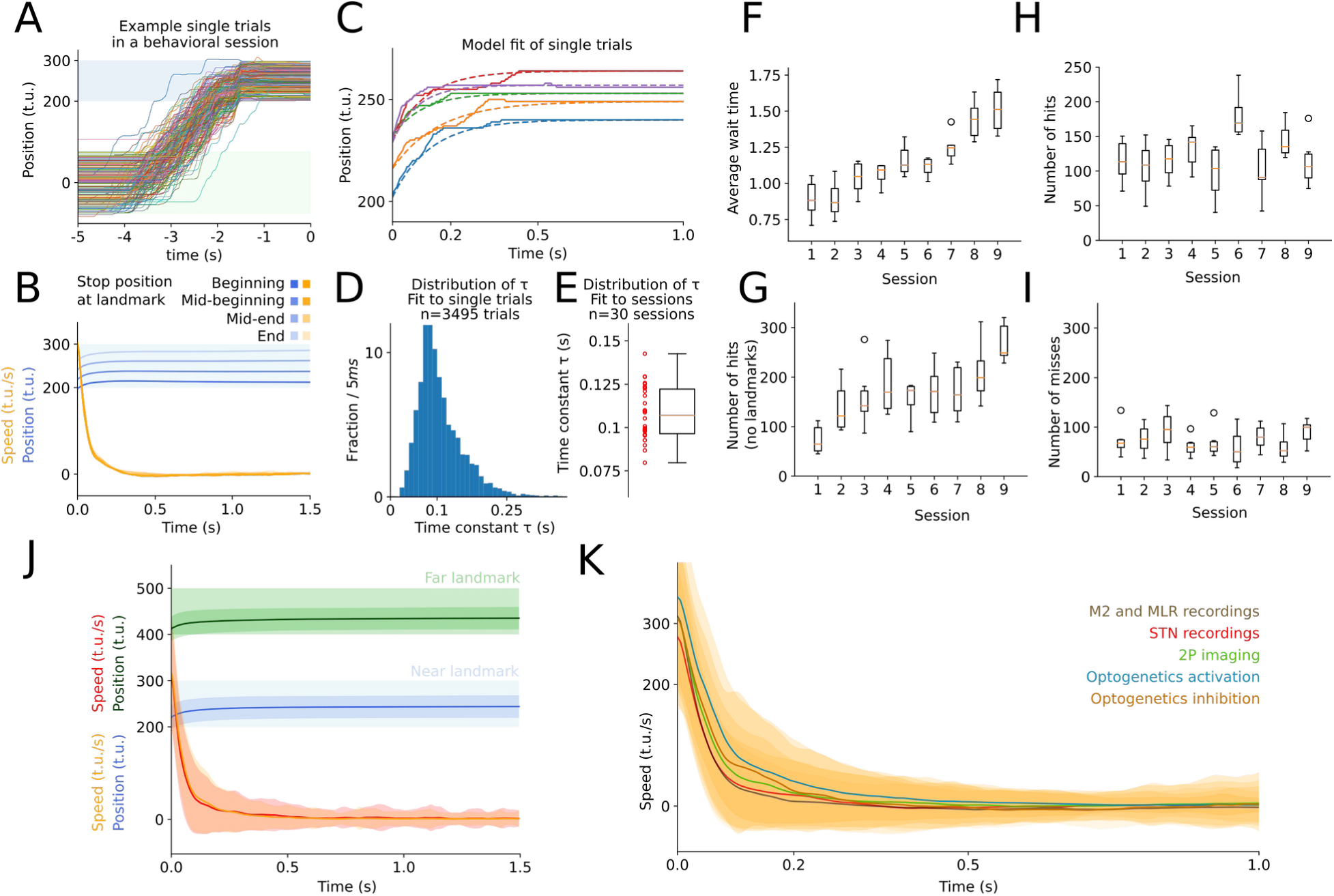
Parameters of the behavioral model and model fits. **(A)** Plot showing example position traces within one behavioral session. The end stopping position at the landmark covers the landmark width. **(B)** Graph showing averages of position and speed of the animal in landmark-stop windows after a switching point. Each average corresponds to one of four intervals of end positions at the landmark. N=10 animals, 3 sessions each. Although the animal stops at different positions along the landmark across trials, the temporal shape of the stopping pattern remains the same. **(C)** Plot showing how the behavioral model can approximate landmark stops at the single trial level. The solid line are the position traces of the animal, and the dashed line are the model fit. **(D)** Plot showing the distribution of τ obtained by fitting single trials. **(E)** Plot showing the distribution of τ obtained by fitting single trials and averaging the values within a session. **(F)** Plot showing the evolution of the average wait time at the landmark during learning as a function of consecutive training sessions (N=6 mice). **(G)** Plot showing the evolution of the average number of hits during learning in a habituation phase as a function of consecutive training sessions (N=6 mice). In this habituation phase, there are no landmarks, and mice are tasked to stop spontaneously after running and collect reward. **(H)** Same as (F) but showing the average number of hits (N=6 mice). **(I)** Same as (F) but showing the average number of misses (N=6 mice). **(J)** Traces showing the velocity (yellow, red) and position (blue, green) of the animal in versions of the task where the landmark is positioned to be near (centered at position 250) and far (centered at position 450). The velocity traces overlap (N=3 mice). **(K)** Traces showing the average velocity during the task in five main experiments (N=3 mice per trace). Both laser ON and laser OFF trials were included in the average for the optogenetics experiments.

**Supplementary Figure 2:**
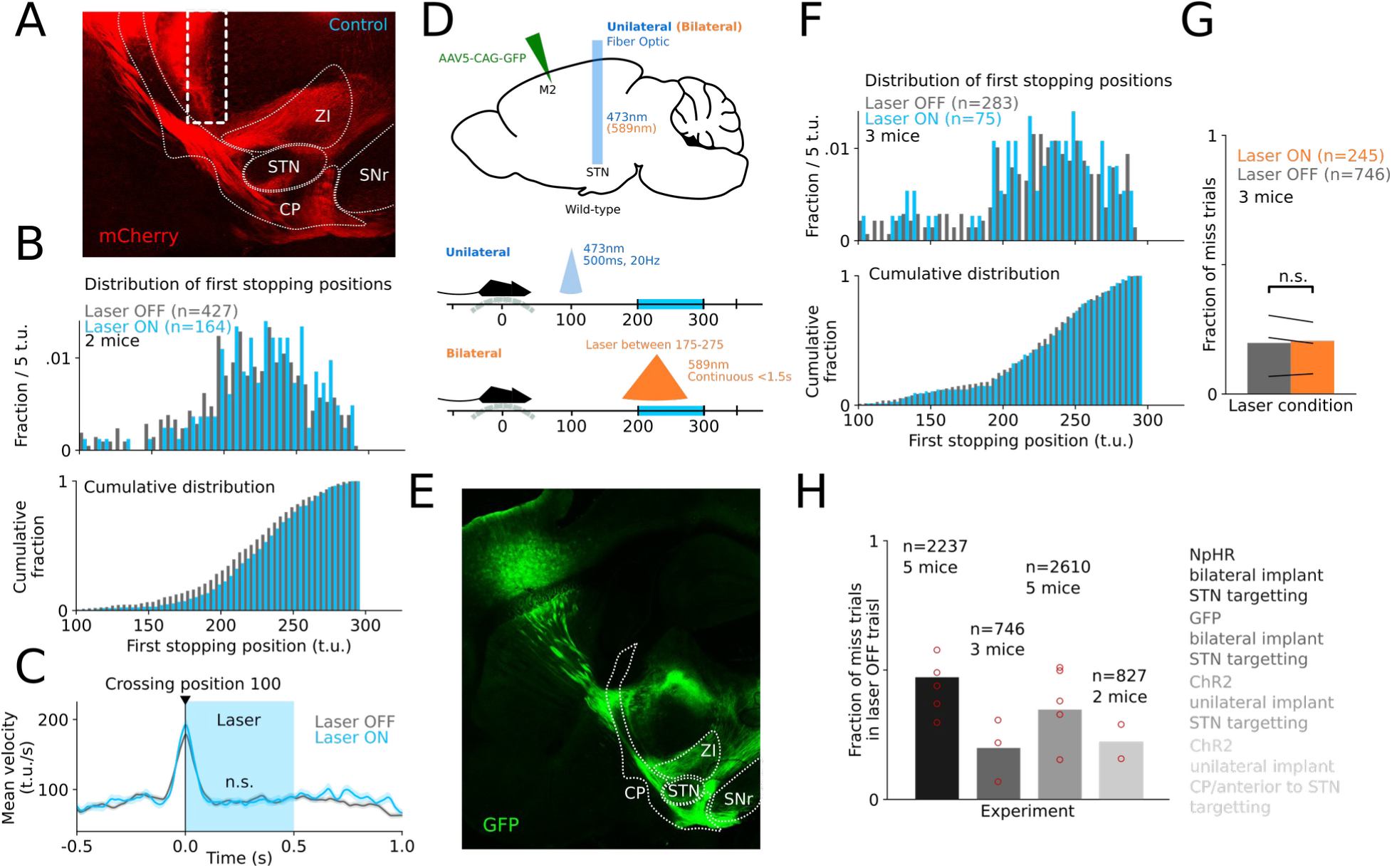
Optogenetics control experiments do not show effects on stopping. **(A)** An AAV virus expressing ChR2 was unilaterally injected in M2 of wild-type mice (N=2) and an optic fiber implanted over the Cerebral Peduncle (ipsilateral to the injection site) anteriorly adjacent to STN to optogenetically target the M2 efferent fiber tract. **(B)** Plots showing the distribution of the first position the animal stops at after position 100. **(C)** Plot showing the average velocity aligned to the onset of optogenetic stimulation of M2 efferent fiber tract at position 100 on the track. The plot does not show a difference in average velocity on laser trials and non-laser trials (Mann–Whitney U test, n.s.). **(D)** Schematic detailing control experiments for the unlateral and bilateral optogenetics experiments (N=3 mice for unilateral activation, and N=3 mice for bilateral inhibition). GFP was expressed in M2 instead of ChR2 and NpHR3.0. **(E)** Sagittal section showing M2-STN projections expressing GFP. **(F)** Plots showing the distribution of the first position the animal stops at after position 100 for the activation control experiment. **(G)** Plot showing the fraction of miss trials during laser ON and OFF trials for the inhibition control experiment. **(H)** Plot showing the fraction of miss trials during laser OFF trials in the different optogenetics experiments and control experiments. The high number of misses in the NpHR axonal inhibition experiment suggests an effect that extends beyond laser ON trials to laser OFF trials.

**Supplementary Figure 3:**
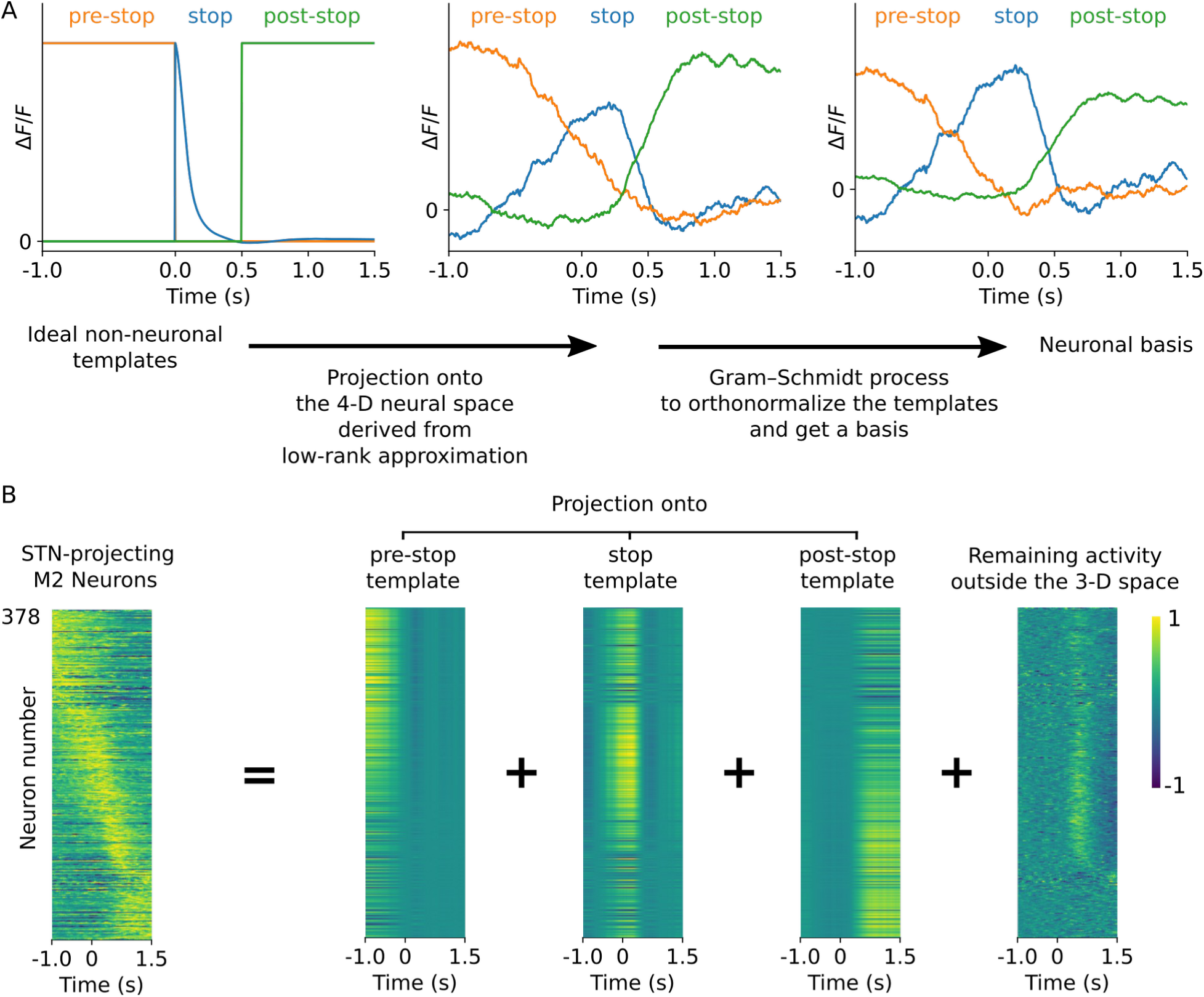
Clustering neuronal activity with dimensionality reduction. **(A) Left.** The three initial non-neuronal templates defined for pre-stop, stop and post-stop activity. **Middle.** The projections of the initial three non-neuronal templates onto a 4-dimensional subspace derived through low-rank approximation. These correspond to the neural responses that represent the non-neuronal templates. **Right.** The orthonormalized responses that form a basis, generating a 3-dimensional subspace, upon which we can decompose our signals. **(B)** Each neuronal response can be written as a combination of four components: the first three correspond to weighted versions of the basis functions, and the fourth corresponds to the activity that remains outside the three-dimensional subspace spanned by the basis. The weights on the templates are used to perform clustering. The remaining activity shows some surge after stopping. The 1-dimensional subspace containing the highest energy in the remaining activity will consist of a neural trajectory with activity concentrated after the stop. When added to the three basis functions (pre-stop, stop and post-stop), the new set will span the 4-dimensional space obtained initially. The decomposition is only used for clustering and not to post-process neural signals.

**Supplementary Figure 4:**
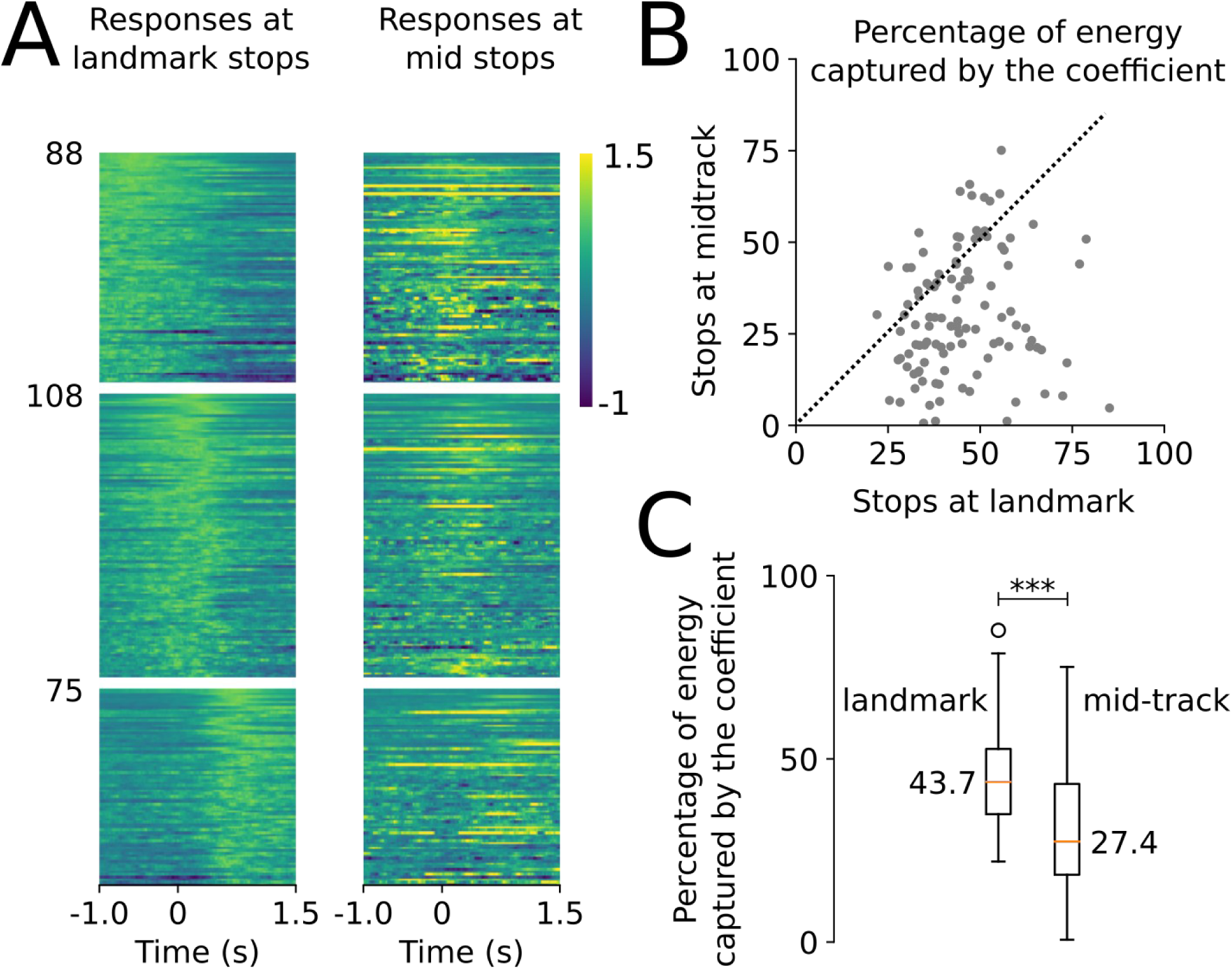
Analysis of landmark and mid-track stops. **(A) Left.** Graph showing the neurons (271 out of 378), whose neural response at landmark stops, had its energy (area under the squared signal) more than 80% explained by the subspace, clustered into three groups (pre-stop, stop and post-stop) using the templates in Figure 4B. **Right.** Graph showing the neural responses of the same neurons, in the same ordering, but on spontaneous stops. The landmark-stop responses are normalized, and the spontaneous stop responses are normalized by the peak value of the landmark-stop responses. **(B)** Graph showing a scatterplot of the percentage of energy captured by the stop template of Figure 4C, in the responses for spontaneous-stops vs landmark-stops. This percentage is the coefficient in Figure 4D divided by the total energy of the signal. Each data point represents a stop neuron (N = 108) taken from (A). **(C)** Graph showing box plots for the distribution of the percentage in (C). The orange lines represent the respective medians. The mean of the distributions is significantly different (Paired t-test, ***:p=1.01e-11).

**Supplementary Figure 5:**
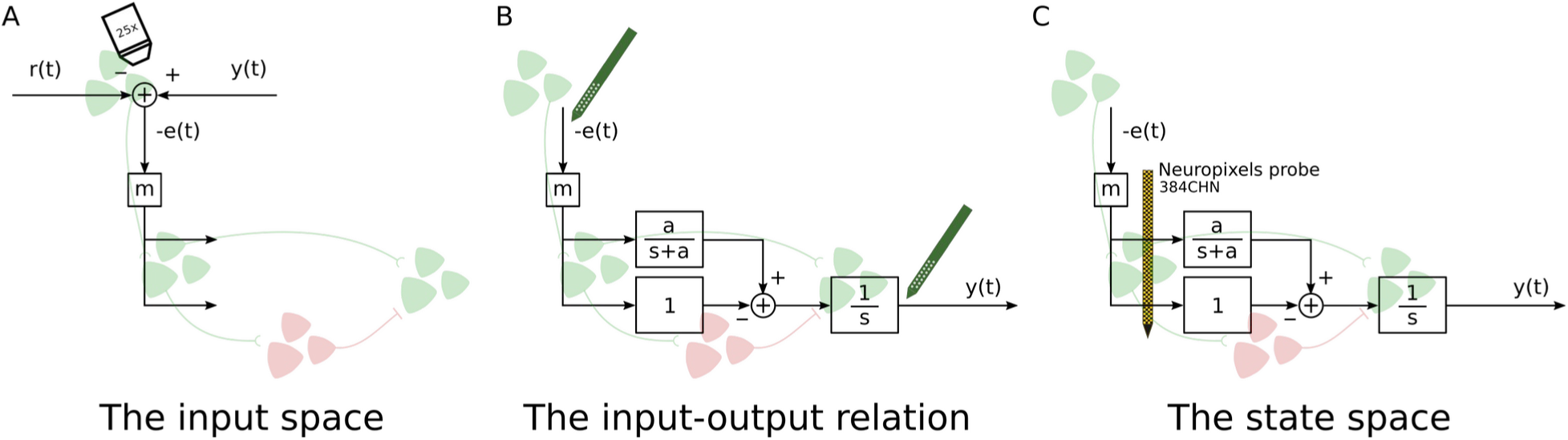
Three key measurements are required to characterize the controller. **(A)** To characterize the input space, we imaged the activity of M2 neurons projecting to STN. **(B)** To characterize the input-output relation, we recorded extracellular single-units simultaneously in M2 and MLR. **(C)** To characterize the dynamical state of the controller, we recorded extracellular single-unit activity in STN.

**Supplementary Figure 6:**
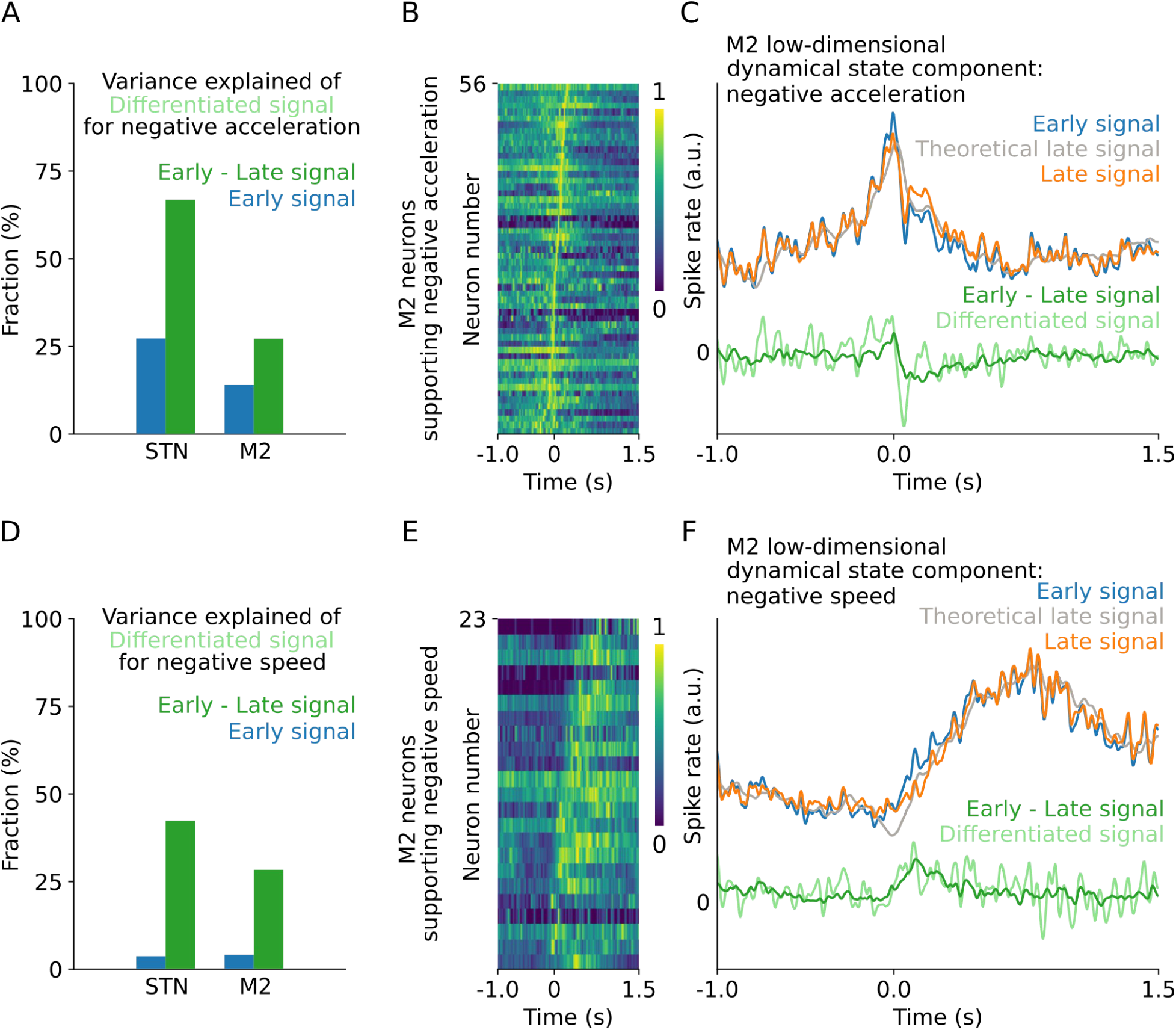
The neural signatures of a dynamical state are absent in M2. **(A)** We repeated the exact analysis performed for STN in Figure 7B-D to examine if a low-dimensional signal could support the negative acceleration component of the dynamical state in M2. The graph shows the explained variance for the STN and M2 population. The variance explained by the M2 population is lower than that explained by the STN population. More importantly, the M2 population cannot recreate the trough (local minimum) following the peak in the differentiated signal incurred by differentiation seen in (C). **(B)** Plot showing M2 neurons whose activity peaked between 250ms before and 250ms after the onset of stops. The neurons are ordered by peak timing. **(C)** Using the population of (B), as performed in Figure 7D, we derived the best two low-dimensional signals representing the negative acceleration component of the error signal (early signal), and its dynamical state counterpart (late signal). **(D)** We repeated the exact analysis performed for STN in Figure 7B-D to find a low-dimensional signal could support the negative speed component of the dynamical state in M2. The graph shows the explained variance for the STN and M2 population. Though the variance for the non-negative speed is lower for M2, the essential difference in explained variance for M2 lies in explaining the negative acceleration component (A-E). **(E)** Plot showing neurons whose activity transitioned from low to high between 250ms before and 750ms after the onset of stops. The neurons are ordered by transition timing. **(F)** Using the population of (E), as performed in Figure 7G, we derived the best two low-dimensional signals representing the negative speed component of the error signal (early signal), and its dynamical state counterpart (late signal).

## Supplementary Text 1: Behavioral model through optimal control

We modeled the behavior of the animal in a single trial as a minimum-time optimal-control problem. The mice were water-regulated, and we assumed that a trained mouse strives to collect the maximum amount of reward possible during the session, while its motivation is sustained. As the trials are of variable duration contingent on the animal’s movement, maximizing reward is equivalent to minimizing time in a trial to collect reward. From an optimal-control perspective, we stated the problem as follows. Starting from an initial position away from the landmark, the mouse is tasked to pick a locomotor plan (control policy) that dictates its locomotion pattern so as to minimize time to collect reward. This time is minimized by minimizing the time needed to reach the landmark position and achieve zero velocity, and thereby initiate waiting at the landmark for 1.5s. The task was based only on positive reinforcement; the animal was not punished for miss trials. Furthermore, the animal was allowed to consider any locomotion trajectory to collect a reward, as long as it held its position at the landmark for 1.5 seconds. As such, there are no major constraints on the locomotor plan, except (a) how it affects the locomotion speed of the animal, and (b) that there is a maximal speed that the animal can achieve. For (a), the locomotion plan (in t.u./s) is a scaled version of the desired trajectory that the speed of the animal should follow. Whenever the speed is lower than the value dictated by the scaled locomotion plan, the speed should increase, and vice versa. We modeled this by having the difference between the plan scaled by a number (αu_t_) and the speed (v_t_) dictate the rate of change in speed, namely the acceleration, up to a multiplicative factor τ. This relation is captured by the simplest possible dynamics as a first-order ordinary differential equation between the speed of the animal v_t_ and the locomotor plan u_t_, parametrized by a time-constant τ:

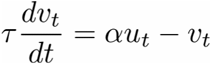

The parameter τ is fixed and dictates how quickly the speed follows the locomotion plan. Specifically, if τ is 0, then the speed is equal to the scaled plan. However, if τ tends to infinity, then the rate of change dv_t_/dt becomes 0, and the speed never changes despite variations in the plan. The parameter α only makes our modelling more general, as it can always be fixed to 1. For (b), as the animal cannot run infinitely fast and thereby has a maximum achievable speed, we let the locomotor plan u_t_ be bounded between two values, u_min_ and u_max_ to ensure this condition. However, the locomotor plan is allowed to take, for each time point, any value between u_min_ and u_max_. We further added boundary conditions dictating the animal’s initial position, and the desired final position and velocity: the position of the landmark and zero, respectively. More formally, the control policy u_t_ (representing the locomotion plan) is chosen to minimize T, the time to go from position d_0_ and arrive at the landmark at position d_T_ with velocity v_T_ = 0, indicating that the animal just halted. The model can be written in the following form:

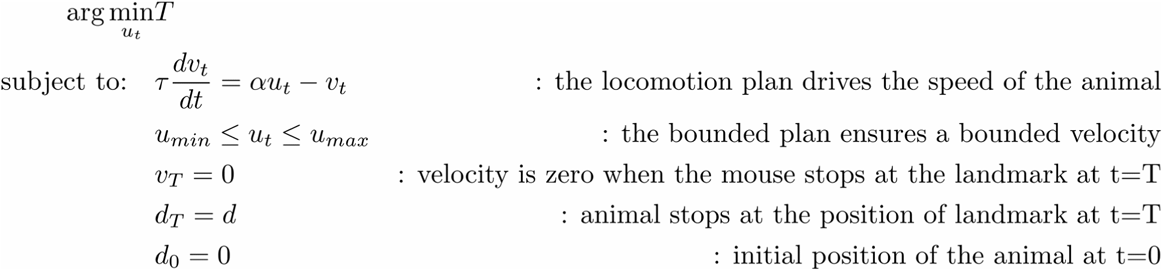

This problem admits an optimal solution for any starting point. The solution can be arrived at by constructing the Hamilton-Jacobi-Bellman (HJB) equation and solving it (Bertsekas, 1995). For a linear system, as is the case here, the optimal solution will be a bang-bang policy of control. As the linear system is first-order, the solution is known to admit at most one switching point (See Example 3.4.3 of (Bertsekas, 1995)). For instance, if d_T_ = 250, namely the landmark is at position 250, then the optimal solution consists of forcing u_t_ equals u_max_ until the animal reaches a switching time point, then u_t_ becomes u_min_ so that the animal stops at position 250.

## Supplementary Text 2: Physiological model and theory

We formalized the physiological dynamics through a feedback control system model (**Figure 5**). The modeled system is a linear time-invariant system (Aström and Murray, 2010; Oppenheim et al., 1996), where the input-output relations (impulse responses) of its various block components are given through the Laplace domain (transfer functions). As convolution in the time domain becomes multiplication in the Laplace domain, composition of systems then consists of multiplying rational functions and cancelling out common factors in the numerator and denominator.

Our diagram consists of a system to control (MLR), a controller (M2-STN) and a negative feedback loop. The MLR was modeled as an ideal integrator, whose transfer function is given by 1/*s*. The controller consists of the difference between the STN-MLR and STN-SNr-MLR, yielding *s*/(*s*+*a*) as a transfer function. Composing the controller with the MLR system amount to multiplying 1/*s* by *s*/(*s*+*a*), which yields 1/(*s*+*a*). This has the effect of substituting the in the denominator of the MLR system by *s*+*a* . The goal of our experiments is to recover through the M2-MLR recordings, and the dynamical state θ of the controller through the STN recordings.

### The input-output relation of the controller-plant system

By computing the closed-loop dynamics, we recovered the effect of the parameter *a* on the decay of MLR activity. In particular, if *R(s)* and *Y(s)* represent the Laplace transforms of *r(t)* and *y(t)* respectively, we then get *Y(s)/R(s)* = *m*/(*s*+*m*+*a*) , yielding an impulse response of *me^−(m+a)t^ u(t)*. If the locomotion plan *r(t)* is at a constant baseline value and then transitions to a new constant lower value and remains there, the decay of *y(t)* will then have a rate of *m+a*. Two terms can then theoretically affect the speed of decay: how much we amplify the error signal in the closed-loop (via *m*) and how much we smooth the differentiated signal (via *a*). It is then theoretically possible to drive the fast decay via only the variable *m*, if we are only investigating the relation between the input *r(t)* and the output *y(t)*. However, by investigating the input-output relation of the controller, we can determine the exact contribution of *a*, and thereby show that amplification is not enough.

Indeed, the open-loop transfer function is given by *Y(s)/E(s)* = *m*/(*s* +*a*), and as such, we have: *e* = (1/*m*)*y* + (*a/m*)*dy/dt*. The error signal is then a weighted combination of two components: a speed signal and an acceleration signal. At the switching point of *r(t)* going from high to low, the error becomes negative. As neural responses are non-negative, we instead define *-e(t)* to be the signal sent down from M2 to STN. We then have:

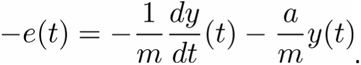

This signal consists of a negative acceleration component and a negative speed component. The m variable consists of only scaling the error signal, and does not interfere with the input-output dynamics of the controller.

### The dynamical state of the controller

If *x(t)* denotes the input to the MLR and *X(s)* its Laplace transform, we have:

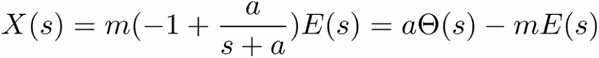

where Θ(*s*) = (1/(*s+a*))*E(s)* denotes the Laplace transform of the dynamical state. In the time-domain, we get the following characterization:

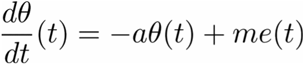

where corresponds to a version of *e(t)* smoothed by an exponential decay function of decay rate *a*. The one-dimensional signal is the dynamical state of the controller. The output of the MLR corresponds to integrating the difference between the dynamical state and the input to STN:

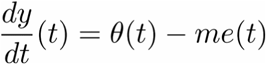

Our model diagram further suggests that differentiation is implemented by the opposing pathway from STN to MLR. In the simplest setting, it specifically suggests that *-me(t)* is provided by the STN-SNr-MLR pathway, whereas *θ(t)* is provided by the STN-MLR projection to MLR. However, other weighted combinations of these signals also yield the same outcomes. Specifically, for non-negative scalars *λ_θ_* and *λ_e_*, we get:

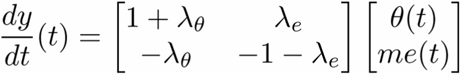

This mathematical characterization is crucial as it indicates that both types of signals can be sent along each pathway and still achieve differentiation. Each signal is only required to be slightly amplified on its corresponding pathway (STN-MLR for *θ* and STN-SNr-MLR for *-me*) as compared to the opposing pathway. The exact weights yielding the contribution of these two signals along a pathway will depend, at least, on parameters pertaining to projection neuron populations. However, the existence of *θ(t)* and *me(t)* are independent of these parameters, and are necessary to be further transmitted from STN to MLR.

## Supplementary Video 1: Example video of a mouse performing the task

The video is submitted along the manuscript.

